# Noise correlations for faster and more robust learning

**DOI:** 10.1101/2020.10.15.341768

**Authors:** Matthew R. Nassar, Daniel Scott, Apoorva Bhandari

## Abstract

Distributed population codes are ubiquitous in the brain and pose a challenge to downstream neurons that must learn an appropriate readout. Here we explore the possibility that this learning problem is simplified through inductive biases implemented by stimulus-independent noise correlations that constrain learning to task-relevant dimensions. We test this idea in a set of neural networks that learn to perform a perceptual discrimination task. Correlations among similarly tuned units were manipulated independently of overall population signal-to-noise ratio in order to test how the format of stored information affects learning. Higher noise correlations among similarly tuned units led to faster and more robust learning, favoring homogenous weights assigned to neurons within a functionally similar pool, and could emerge through Hebbian learning. When multiple discriminations were learned simultaneously, noise correlations across relevant feature dimensions sped learning whereas those across irrelevant feature dimensions slowed it. Our results complement existing theory on noise correlations by demonstrating that when such correlations are produced without significant degradation of the signal-to-noise ratio, they can improve the speed of readout learning by constraining it to appropriate dimensions.

**Significance statement:** Positive noise correlations between similarly tuned neurons theoretically reduce the representational capacity of the brain, yet they are commonly observed, emerge dynamically in complex tasks, and persist even in well-trained animals. Here we show that such correlations, when embedded in a neural population with a fixed signal to noise ratio, can improve the speed and robustness with which an appropriate readout is learned. In a simple discrimination task such correlations can emerge naturally through Hebbian learning. In more complex tasks that require multiple discriminations, correlations between neurons that similarly encode the task-relevant feature improve learning by constraining it to the appropriate task dimension.

## Introduction

The brain represents information using distributed population codes in which particular feature values are encoded by large numbers of neurons. One advantage of such codes is that a pooled readout across many neurons can effectively reduce the impact of stimulus-independent variability (noise) in the firing of individual neurons (Pouget et al., 2000). However, the extent to which this benefit can be employed in practice is constrained by noise correlations, or the degree to which stimulus-independent variability is shared across neurons in the population (Averbeck et al., 2006). In particular, positive noise correlations between neurons that share the same stimulus tuning can reduce the amount of decodable information in the neural population (Averbeck et al, 2006; Moreno-Bote et al., 2014; Hu et al., 2014). Despite their detrimental effect on encoding, noise correlations of this type are reliably observed, even after years of training on perceptual tasks (Cohen and Kohn, 2011). Furthermore, noise correlations between neurons are dynamically enhanced under conditions where two neurons provide evidence for the same response in a perceptual categorization task (Cohen and Newsome, 2008), raising questions about whether they might serve a function rather than simply reflecting a suboptimal encoding strategy.

At the same time, learning to effectively read out a distributed code also poses a significant challenge. Learning the appropriate weights for potentially tens of thousands of neurons in a low signal-to-noise regime is a difficult, high-dimensional problem, requiring a very large number of learning trials and entailing considerable risk of “over fitting” to specific patterns of noise encountered during learning trials. Nonetheless, people and animals can rapidly learn to perform perceptual discrimination tasks, albeit with performance that does not approach theoretically achievable levels (Hawkey et al., 2004; Stringer et al., 2019). In comparison, deep neural networks capable of achieving human level performance typically require a far greater number of learning trials than would be required by humans and other animals (Tsividis et al., 2017). This raises the question of how brains might implement inductive biases to enable efficient learning in high dimensional spaces.

Here we address open questions about noise correlations and learning by considering the possibility that noise correlations facilitate faster learning. Specifically, we propose that noise correlations aligned to task relevant dimensions could reduce the effective dimensionality of learning problems, thereby making them easier to solve. For example, perceptual stimuli often contain a large number of features that may be irrelevant to a given categorization. At the level of a neural population, individual neurons may differ in the degree to which they encode task irrelevant information, thus making the learning problem more difficult. In principle, noise correlations in the relevant dimension could reduce the effects of this variability on learned readout. Such an explanation would be consistent with computational analyses of Hebbian learning rules (Oja, 1982), which can both facilitate faster and more robust learning (Krotov and Hopfield, 2019), and in turn may induce noise correlations. We propose that faster learning of an approximate readout is made possible through low dimensional representations that share both signal and noise across a large neural population. In particular, we hypothesize that representations characterized by enhanced noise correlations among similarly tuned neurons can improve learning by focusing adjustments of the readout onto task relevant dimensions.

We explore this possibility using neural network models of a two-alternative forced choice perceptual discrimination task in which the correlation among similarly tuned neurons can be manipulated independently of the overall population signal-to-noise ratio. Within this framework, noise correlations, which can be learned through Hebbian mechanisms, speed learning by forcing learned weights to be similar across pools of similarly tuned neurons, thereby ensuring learning occurs over the most task relevant dimension. We extend our framework to a cued multidimensional discrimination task and show that dynamic noise correlations similar to those observed in vivo (Cohen and Newsome, 2008), speed learning by constraining weight updates to the relevant feature space. Our results demonstrate that when information is extrinsically limited, noise correlations can make learning faster and more robust by controlling the dimensions over which learning occurs.

## Materials and Methods

Our goal was to understand the computational principles through which correlations in the activity of similarly tuned neurons affect the speed with which downstream neurons could learn an effective readout. Previous work has demonstrated that manipulating noise correlations while maintaining a fixed variance in the firing rates of individual neurons leads to changes in the theoretical encoding capacity of a neural population (Averbeck et al., 2006; Moreno-Bote et al., 2014). To minimize the potential impact of such encoding differences, we took a different approach; rather than setting the variance of individual neurons in our population to a fixed value, we set the signal-to-noise ratio of our population to a fixed value. Thus, our approach does not ask how maximum information can be packed into a given neural population’s activity, but rather how the strategy for packing a *fixed* amount of information in a population affects the speed with which an appropriate readout of that information can be learned. We implement this approach in a set of neural networks described in more detail below.

### Learning readout in perceptual learning task

Simulations and analyses for a simple perceptual discrimination task were performed with a simplified and statistically tractable two-layer feed-forward neural network (figure 3A). The input layer consisted of two homogenous pools of 100 units that were each identically “tuned” to one of two motion directions (left, right). On each trial normalized firing rates for the neural population were drawn from a multivariate normal distribution that was specified by a vector of stimulus-dependent mean firing rates (signal: +1 for preferred stimulus, −1 for non-preferred stimulus) and a covariance matrix. All elements of the covariance matrix corresponding to covariance between units that were “tuned” to different stimuli were set to zero. The key manipulation was to systematically vary the magnitude of diagonal covariance components (eg. noise in the firing of individual units) and the “same pool” covariance elements (eg. shared noise across identically tuned neurons) while maintaining a fixed level of variance in the summed population response for each pool:

**Figure 1:**
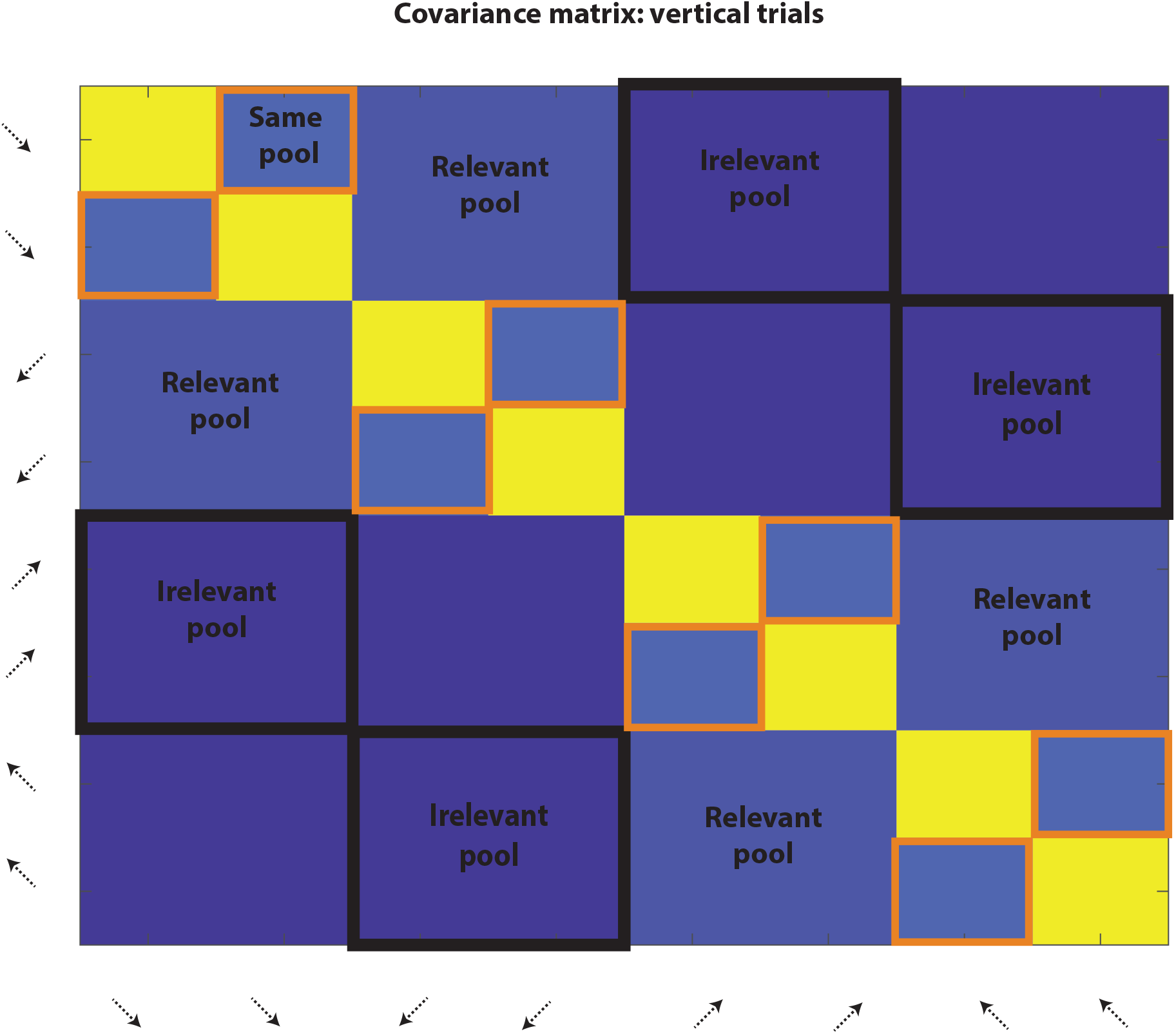
Schematic of covariance matrix for two-dimensional motion discrimination task. The covariance between units with different motion tuning (reflected by the arrows labeling columns and rows) is schematically represented for a simplified input layer, where only two identically tuned neurons are in each pool (in actual simulations there were 100 units per pool). Same pool correlations are controlled by covariance elements between neurons with identical tuning (orange boxes). Relevant pool correlations are controlled by covariance elements between neurons that are similarly tuned to the task-relevant feature. Task irrelevant correlations are controlled by covariance elements between neurons that are similarly tuned to the task-irrelevant feature. The covariance matrix shown here is for a vertical trial – on a horizontal trial the irrelevant pool and relevant pool locations would be reversed. Covariance elements for pairs of neurons that differed in tuning on both dimensions were set to zero. Each input population has been depicted as two units here for presentation purposes. Background color reflects the case where same pool correlations = 0.2 and relevant pool correlations = 0.1.

**Figure 2:**
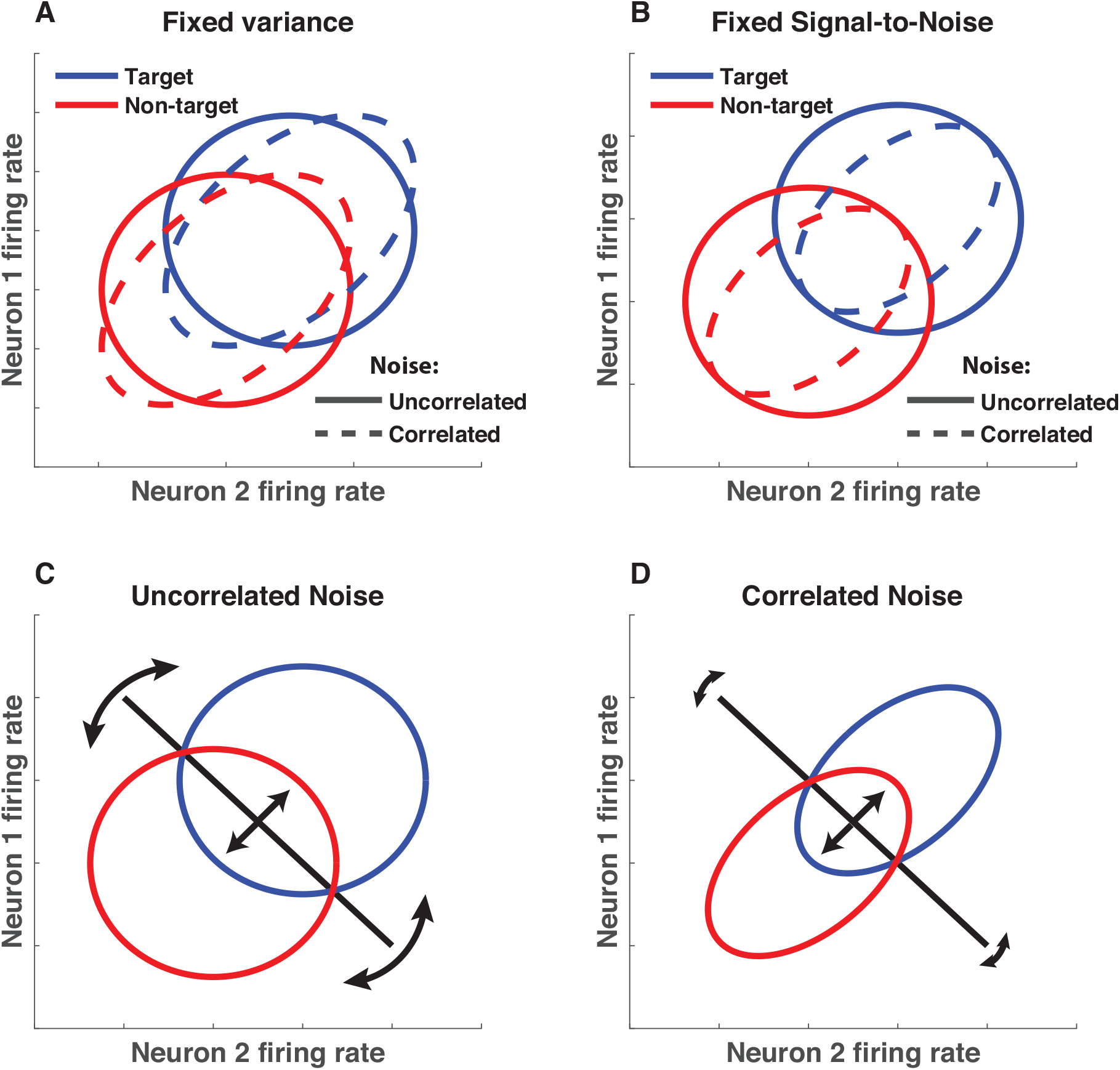
Modeling noise correlations with extrinsic constraint on signal-to-noise ratio. **A)** Previous work has modeled noise correlations by assuming that population variance is fixed and that covariance is manipulated to produce noise correlations. Under such assumptions, the firing rate of two similarly tuned neurons is plotted in the absence (solid) or presence (dotted) of information-limiting noise correlations. **B)** Here we assume that the signal-to-noise ratio of the neural population is limited to a fixed value such that noise correlations between similarly tuned neurons do not affect theoretical performance. Thus, the percent overlap of blue (target) and red (non-target) activity profiles does not differ in the presence (dotted) or absence (solid) of noise correlations. **C&D)** Under this assumption, noise correlations among similarly tuned neurons could compress the population activity to a plane orthogonal to the optimal decision boundary, thereby minimizing boundary adjustments in irrelevant dimensions (**C**) and maximizing boundary adjustments on relevant ones (**D**).

**Figure 3:**
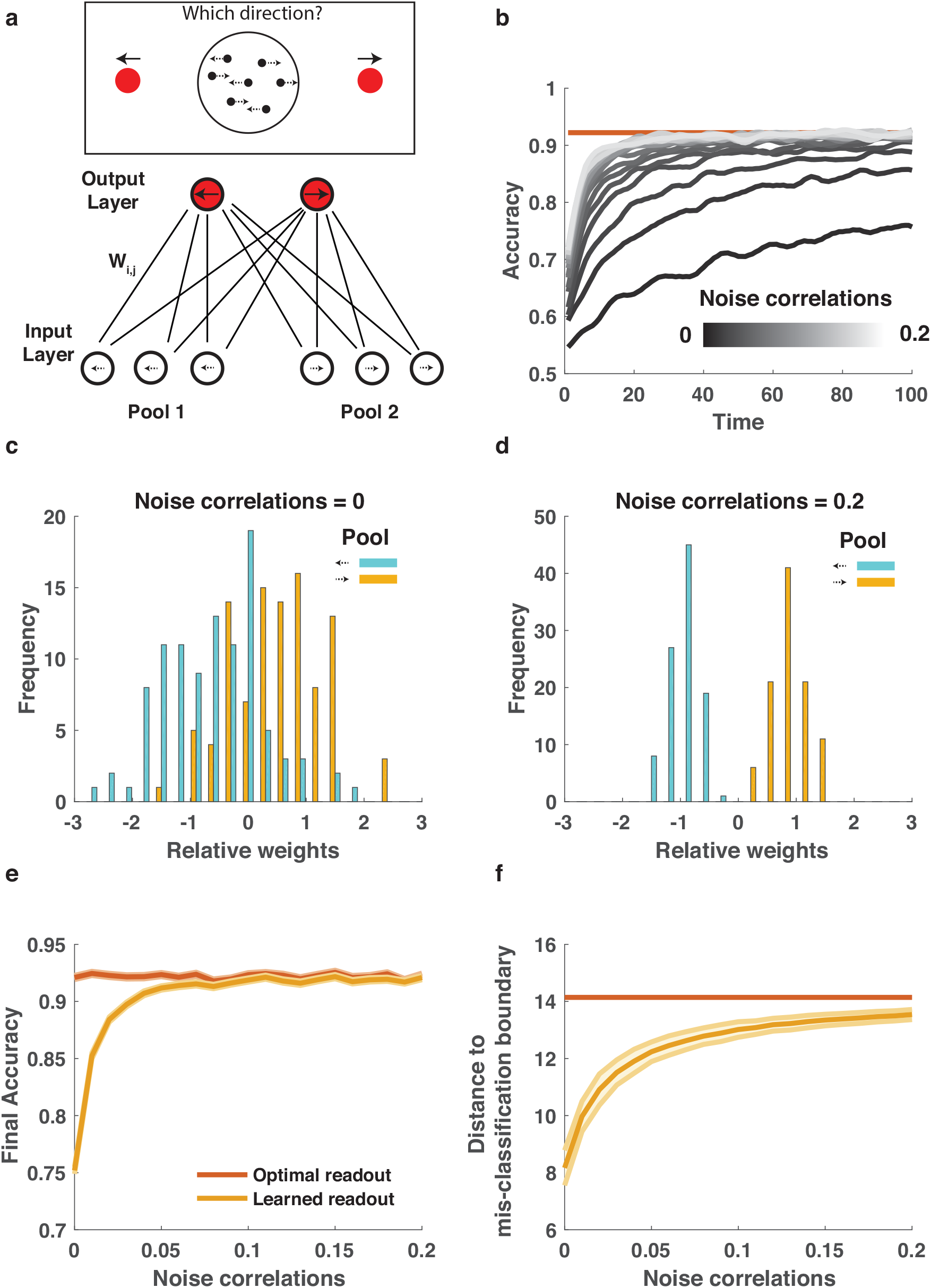
Correlated noise within similarly tuned populations leads to faster and more robust learning of a perceptual discrimination. **A)** A two-layer feed-forward neural network was designed to solve a two alternative forced choice motion discrimination task at or near perceptual threshold. Input layer contains two homogenous pools of identically tuned neurons that provide evidence for alternate percepts (eg. leftward motion versus rightward motion) and output neurons encode alternate courses of actions (eg. saccade left versus saccade right). Layers are fully connected with weights randomized to small values and adjusted after each trial according to rewards (see methods). **B)** Average learning curves for neural network models in which population signal-to-noise ratio in pools 1 and 2 were fixed, but noise correlations (grayscale) were allowed to vary from small (dark) to large (light) values. **C&D)** Weight differences (left output – right output) for each input unit (color coded according to pool) after 100 timesteps of learning for low (**C**) and high (**D**) noise correlations. **E**) Accuracy in the last 20 training trials is plotted as a function of noise correlations for learned readouts (orange) and optimal readout (red). Lines/shading reflect Mean/SEM. F) The shortest distance, in terms of neural activation, required to take the mean input for a given category (eg. left or right) to the boundary that would result in misclassification is plotted for the final learned (orange) and optimal (red) weights for each noise correlation condition (abscissa). Lines/shading reflect Mean/SEM.

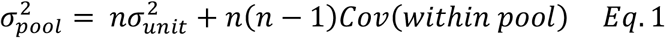

Where 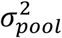 is the variance on the sum of normalized firing rates from neurons within a given pool, n is the number of units in the pool and the within pool covariance (*Cov*(*within pool*)) specifies the covariance of pairs of units belonging to the same pool. Signal-to-noise ratio (SNR) was defined as the population signal (preferred-antipreferred) divided by the standard deviation of the population response in the signal dimension. SNR was set to be 2 for each individual pool of neurons, leading to a signal-to-noise ratio for the entire population (both pools) equal to 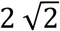. Given this constraint, the fraction of noise that was shared across neurons within the same pool was manipulated as follows:

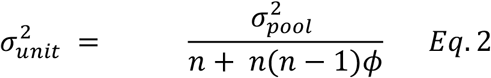

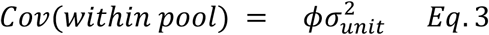

Where *ϕ* reflects the fraction of noise that is correlated across units, which we refer to in the text as noise correlations. Noise correlations (*ϕ*) were manipulated across values ranging from 0 to 0.2 for simulations. Note that, since *ϕ* appears in the denominator of equation 2, adding noise correlations while sustaining a fixed population signal-to-noise ratio leads to lower variance in the firing rates of single neurons, differing from previous theoretical assumptions (compare figure 2a&b).

The input layer of the neural network was fully connected to an output layer composed of two output units representing left and right responses. Output units were activated on a given trial according to a weighted function of their inputs:

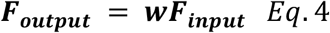

Where *F*_*output*_ is a vector of firing rates of output units, *F*_*input*_ is a vector of firing rates of the input units, and w is the weight matrix. Firing of an individual output unit can also be written as a weighted sum over input unit activity:

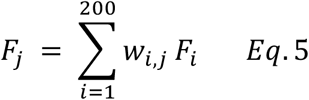

where *F*_*j*_ reflects the firing of the j^th^ output unit, *F*_*i*_ reflects the firing of the i^th^ input unit, and *w*_*i*_,_*j*_ reflects the weight of the connection between the i^th^ input unit and the j^th^ output unit. Actions were selected as a softmax function of output firing rates:

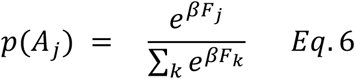

where *β* is an inverse temperature, which was set to a relatively deterministic value (10000). Learning was implemented through reinforcement of weights to the selected output neuron (subscripted j below):

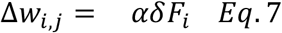

Where *F*_*i*_ is the normalized firing rate of the i^th^ input neuron, *δ* is the reward prediction error experienced on a given trial [+0.5 for correct trials and −0.5 for error trials], and *α* is a learning rate (set to 0.0001 for simulations in figure 2). The network was trained to correctly identify two stimuli (each of which was preferred by a single pool of input neurons) over 100 trials (the last 20 trials of which were considered testing). Simulations were repeated 1000 times for each level of *ϕ* and performance measures were averaged across all repetitions. Mean accuracy per trial across all simulations was convolved with a Gaussian kernel (standard deviation = 0.5 trials) for plotting in figure 2b. Mean accuracy across the final 20 trials was used as a measure of final accuracy (figure 2e). Statistics on model performance were computed as Pearson correlations between noise correlations *ϕ* and performance measures across all simulations and repetitions.

### Analytical learning trajectories

One advantage of our simple network architecture is its mathematical tractability. To complement the simulations described above, we also explored learning in the network analytically. Specifically, we decomposed weight updates into two categories: weight updates in the signal dimension, and weight updates perpendicular to the signal dimension. Weight updates in the signal dimension improved performance through alignment with the signal itself, whereas weight updates in the perpendicular dimension limited performance through chance alignment with trial-to-trial noise. An intuition for our approach and derivation are provided below.

The two-alternative discrimination task is a one dimensional signal detection problem, because it depends only on the difference between two scalars. In particular, if *y* = [*y*_1_, *y*_2_] denotes the readout activity the pair of pools and r denotes the response (e.g. r=-1 is “respond left” and r=1 is “respond right”), then *r* = *r*(*y*_1_ – *y*_2_) = *r*(*Δy*). In addition, *Δy* = *w*_1_*x* − *w*_2_*x* ≡ *Δwx*, where x reflects the firing rates of the input units and w_1_ reflects the vector of weights mapping input activation onto output unit 1 (y_1_). To determine how accuracy is impacted by noise correlations, we ask how Mahalanobis distance (d’), mean separation (d), and signal variance 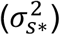 diverge over training time for the different noise correlation conditions. The effective variance, 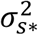, differs from the true noise variance in the signal dimension due to the fact that out-of-signal-dimension noise is transferred into the signal dimension by imperfect readout weights. Intuitively, learning speed may be improved by noise correlations because less out-of-dimension noise is “learned into” the weights, thereby reducing the transfer out-of-dimension noise into the signal space on any given trial.

The logic of training is as follows: On a correct trial, the weights to the chosen unit are incremented by a multiple of the input vector x. That is:

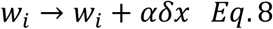

Here α reflects a positive learning rate, x reflects the activity of the input units and δ is the reward prediction error, which we use as the absolute reward prediction error instead of the signed one in this section for convenience.

Now the input is a sum of signal and zero mean noise:

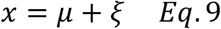

The expectation of noise is zero (E(ξ) = 0) and the signal μ can take only two values μ ∈ {±μ_0_}. Therefore if the weights start from some value Δw(0), we will find that:

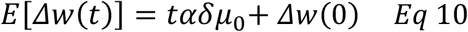

Where t reflects the current timestep of learning. In words, we expect the amount of signal in the weights to increase linearly over time. This means that we expect the response to a noise-free signal (μ_0_) after t timesteps to be:

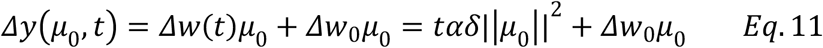

This is the measure d between the two Gaussian peaks in the one dimensional signal detection problem described above. Below, we ignore the initial weight term, since it does not change over time. To compute accuracy and d’ over training time we also need to compute the effective variance along the signal dimension. First we note that the noise can be decomposed as:

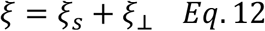

where ξ_s_ and ξ_⊥_ are orthogonal components of the noise in the signal dimension (ξ_s_) and perpendicular to the signal dimension (ξ_⊥_). Here we consider cases where the noise along the signal dimension (ξ_s_) has constant variance, following on the assumptions that SNR is set to a constant value and that the mean signal is the same for all noise correlation conditions.

The difference Δy on any given trial decomposes into a sum of terms, one reflecting weight-based transfer of signal and one reflecting the transfer of orthogonal noise. This latter term arises because the weights are not, at any finite time, a perfect matched filter for the signal. Letting subscripts s and ⊥ continue to denote “signal” and “perpendicular” dimensions, we have:

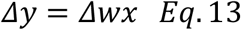

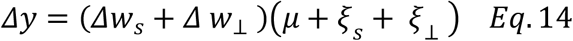

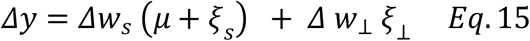

where the final equation reflects the absence of terms that have zero products by definition of the perpendicular subspaces. The variance of Δy can be computed using independence and orthogonality properties:

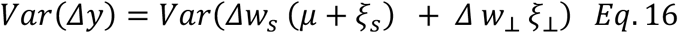

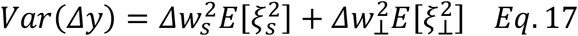

For any given network, the term 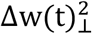 is a mean-zero diffusion process arising from the fact that noise is added to the weights at every time step. For the Gaussian white noise case, 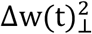 is equivalent to Brownian motion in the (n-1) dimensions perpendicular to the signal. Because (n-1) is not small, the summed empirical variance of these processes, operative on each component, is likely to be close to the theoretical total variance. If we split the term 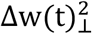 into the (n-1) components and index them with i, this gives:

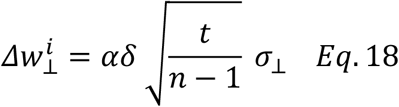

The denominator of (n-1) appears here because Brownian motion determines growth in the variance of each of the (n-1) perpendicular noise directions among which the total variance σ_⊥_is distributed. Technically, our manipulation of the noise covariance fixes the variance in a second direction of the space as well, so that noise variance is actually evenly distributed over only (n-2) of the (n-1) perpendicular dimensions, but this inhomogeneity is inconsequential if n is not small; In effect, we are ignoring an order 1 term relative to an order n term for simplicity. In order to understand how perpendicular weights grow with time, we need only to determine σ_⊥_(ϕ), where ϕ is the parameter controlling the noise covariance matrix in our simulations. Specifically, the first row of the covariance matrix takes the form:

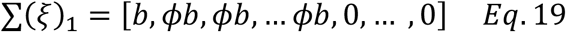

Using the additional fact that row-sums are set to 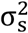 to control the signal variance, we find that:

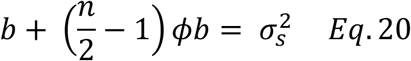

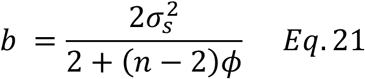

Since the eigenvalues of ∑(ξ) are the variances in different dimensions of the space, we can find the total variance perpendicular to the signal by subtracting the known signal variance from the trace of ∑(ξ):

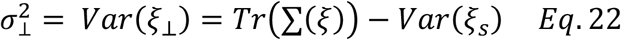

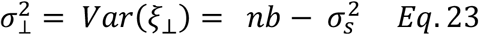

Putting this together with previous results, we have:

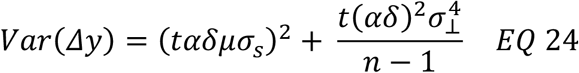

This provides analytic prediction for the variance of our readout decision variable Δy after learning for t trials, using a learning rate α to learn from from prediction errors of magnitude δ. Note that σ_s_ was fixed in our simulations, but that 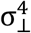 depends on ϕ through b, such that larger values of ϕ lead to smaller values of b, and thus a smaller 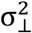, reducing the second term in Eq 24. Furthermore, since the first term in Eq 24 scales with t^2^ its contributions dominate as more trials are observed. This leads to identical asymptotic variance in the limit of large t, since the first term does not depend on ϕ.

By combining the mean and variance information in Equations 11 and 24 we computed accuracy as one minus the cumulative probability density of the Gaussian distribution 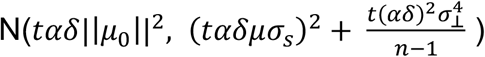 evaluated from negative infinity to zero.

### Noise correlations with fixed signal-to-noise ratio and single unit variance

Noise correlations produced by the simulations above lead to reductions in the overall variance of single unit firing rates. In order to validate that our results depend on maintaining signal-to-noise, rather than depending on single-unit variance, we also consider the case where noise correlations are introduced with a fixed level of single unit variance. In this case, signal-to-noise ratio was maintained by scaling the amount of signal according to the level of noise correlations (see https://github.com/NassarLab/NoiseCorrelation for full derivation):

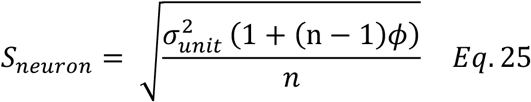

where *S*_*neuron*_ reflects the amount of signal provided by each unit, 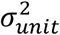 reflects a fixed variance assigned to each unit, n reflects the number of units in the pool, and *ϕ* reflects the level of noise correlations. Thus, when we simulated correlated noise using this equation, neurons maintained the same variance 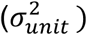, but increased their signal relative to the zero noise correlation condition (*ϕ* = 0).

### Noise correlations that are bounded to a maximum signal-to-noise ratio

In order to examine the importance of our assumption regarding fixed signal-to-noise ratio, we also considered a parameterized model where signal (*S*_*neuron*_) was set according to a linear mixture:

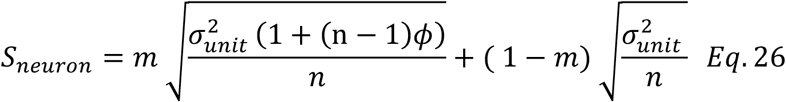

where m is a mixing parameter that combines the signal producing a fixed signal-to-noise ratio (first term) with a fixed signal that does not depend on the level of noise correlations (second term). When m is set to 1, this parameterized model obeys our assumptions regarding fixed signal to noise ratio, but when m is set to 0, the model conforms to more standard assumptions regarding fixed single unit variance and signal.

### Hebbian learning of noise correlations in three-layer network

We extended the two-layer feed-forward architecture described above to include a third hidden layer in order to test whether Hebbian learning could facilitate production of noise correlations among similarly tuned neurons (figure 5A). The input layer was fully connected to the hidden layer, and each layer contained 200 neurons. In the input layer, neurons were homogenously tuned (100 leftward, 100 rightward) as described above, with *ϕ* set to zero (eg. no noise correlations). Weights to the hidden layer were initialized to favor one-to-one connections between input layer units and hidden layer units by adding a small normal random weight perturbation (mean=0, standard deviation = 0.01) to an identity matrix (though an alternate initialization was used to produce figure 6-1). During learning, weights between the input and hidden layer were adjusted according to a normalized Hebbian learning rule:

**Figure 4:**
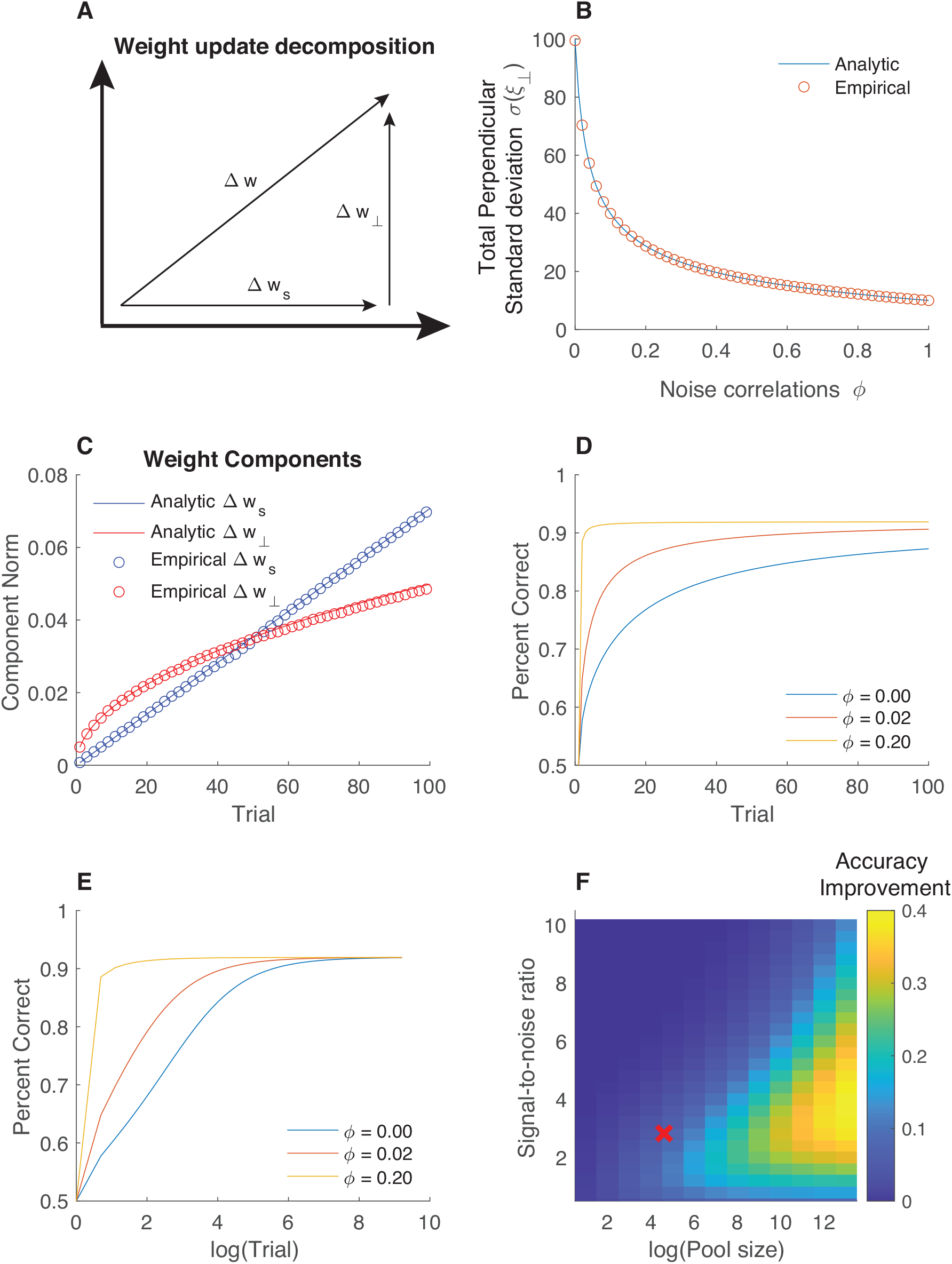
Analytic learning trajectories demonstrate advantage for noise correlations when pools are large and signal-to-noise ratio is low. **A)** Our analytical approach decomposed weight updates *Δw* into two components: updates in the signal dimension (*Δ w*_s_) and updates perpendicular to the signal dimension (Δ w_⊥_). **B**) Standard deviation of the variability in the dimension perpendicular to the signal (ordinate) decreased as a function of noise correlation (abscissa) as derived with our analytic approach (blue line, see methods), and for the empirical simulations. **C)** For a given noise correlation (0.02 in this example) learning yielded weight changes in the signal dimension (blue circles) as well in the perpendicular dimension (red circles) that could be described analytically (blue and red lines). Circles represent average values from twenty empirical simulations. **D&E)** Theoretical accuracy derived from the analytical weights reproduces learning advantages observed in our simulations for higher levels of noise correlations (compare yellow to blue curves) and demonstrates convergence with sufficient observations (**E**; note abscissa in log units). **F)** Improvement in average accuracy over first 100 trials, derived analytically by taking the mean difference between yellow and blue curves in (**D**), is indicated in color across a range of signal-to-noise ratios (ordinate) and neural population sizes (abscissa). The largest learning advantages for noise correlations were observed in large neural populations that contained limited stimulus information (moderately low SNR). Red X depicts parameters used for our simulations.

**Figure 5:**
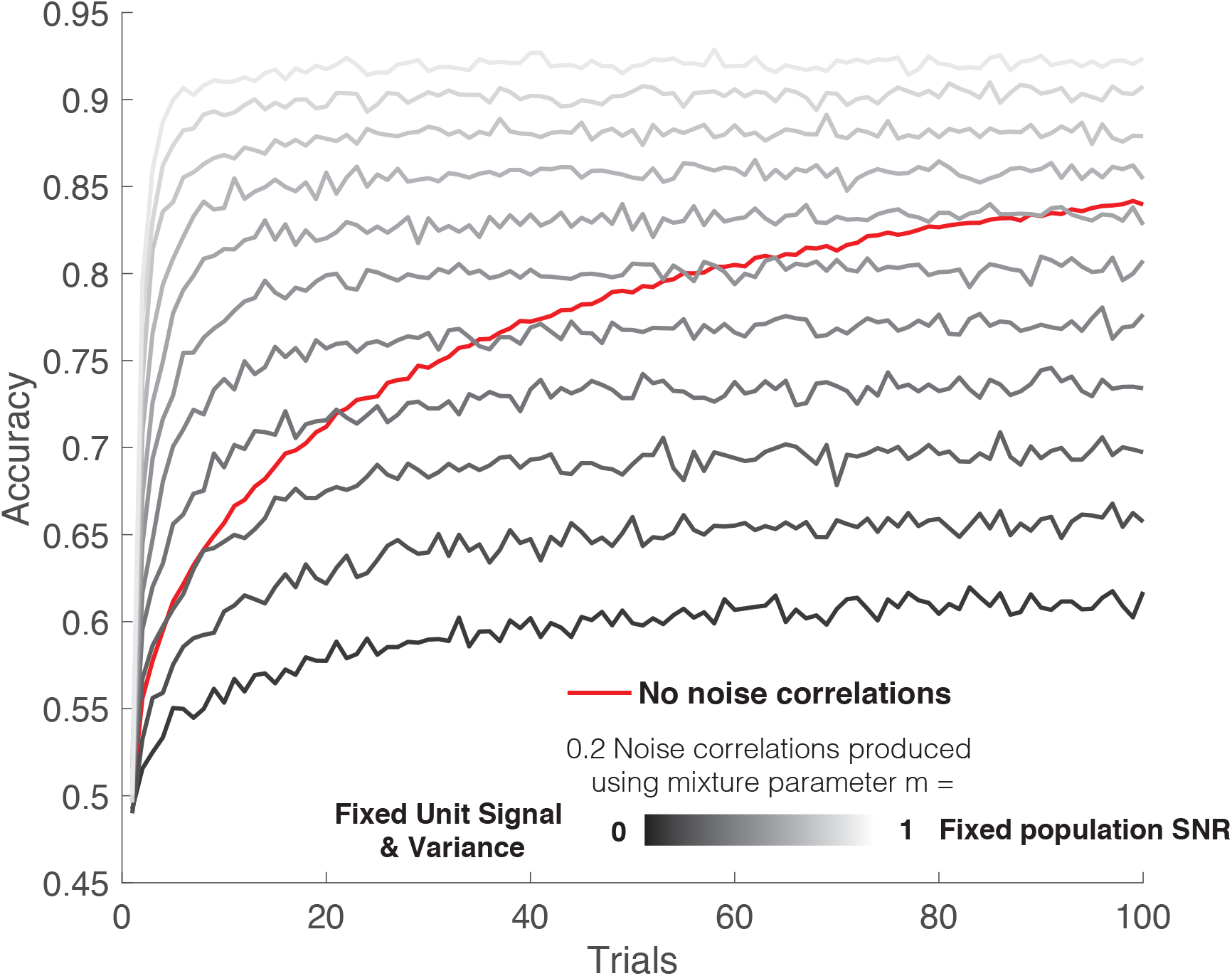
Impact of noise correlations on learning depends on the assumption that signal-to-noise ratio is fixed. Accuracy (ordinate) as a function of trials (abscissa) for a model without noise correlations (red) and for several models that generate noise correlations (0.2) under different assumptions. The lightest color reflects a case where signal-to-noise ratio of the population is completely preserved, analogous to our previous simulations. The darkest color reflects a case where the variance and signal of individual neurons is fixed, leading to a population signal-to-noise ratio that varies as a function of noise correlations. Intermediate colors indicate parametric mixtures of these assumptions created using equation 26. Note that learning advantages depend critically on assumptions about signal-to-noise ratio, and that noise correlations implemented using intermediate assumptions introduce a tradeoff between faster learning (gray lines above red line for early trials) and lower asymptotic performance (gray lines below red line for later trials).

**Figure 6:**
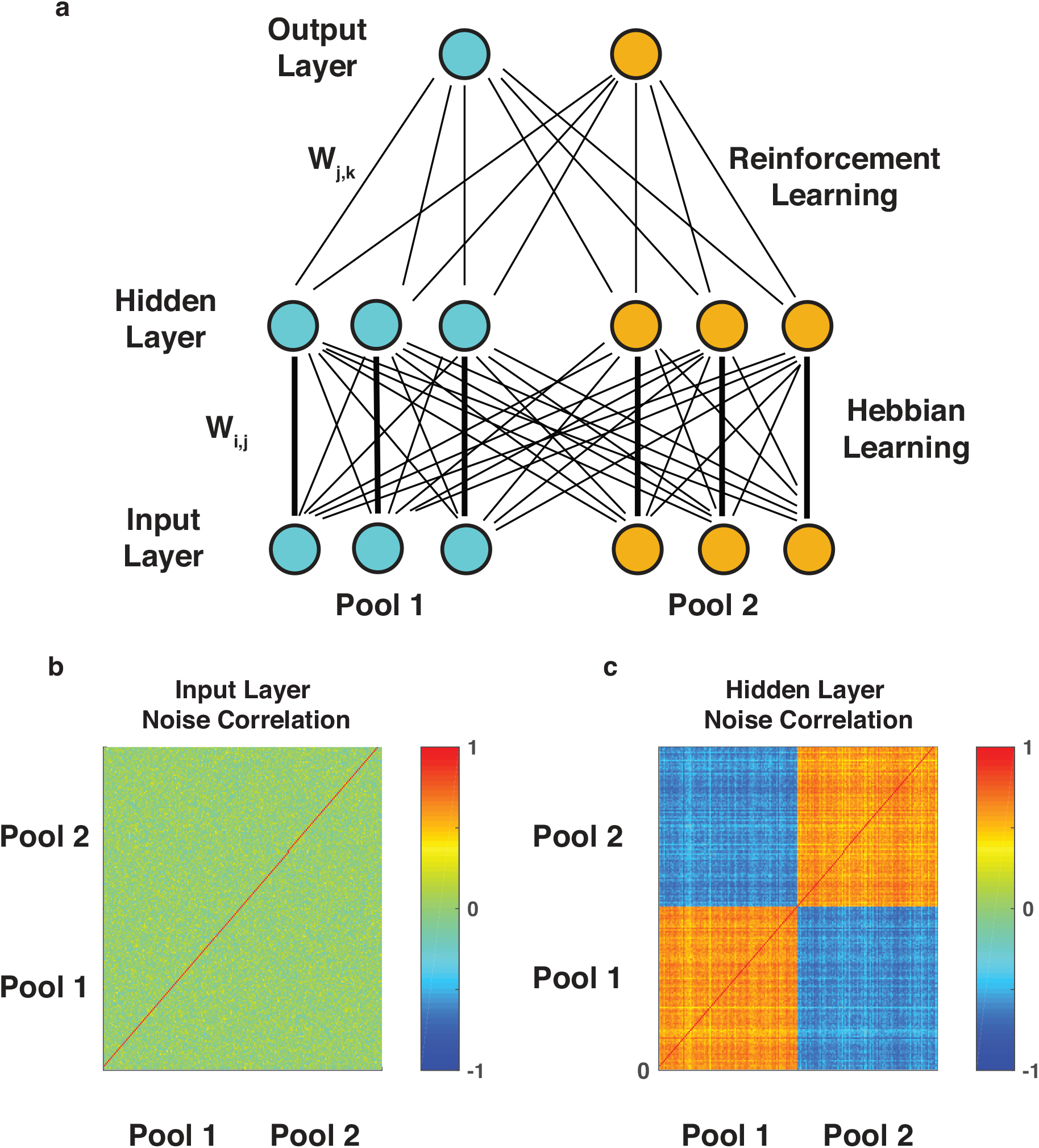
Hebbian learning produces correlations within similarly tuned populations in a perceptual discrimination task. **A**) Three-layer neural network architecture. Input layer feeds forward to hidden layer, which is fully connected to an output layer. Input layer provides uncorrelated inputs to hidden layer through projection weights that are adjusted according to a Hebbian learning rule. **B&C**) Noise correlations observed in hidden layer units at the beginning (**B**) and end (**C**) of training.

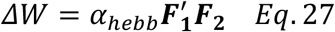

Where 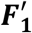 is a normalized vector of firing rates corresponding to the input layer and ***F***_***2***_ is a normalized vector of firing rates corresponding to the hidden layer units. The learning rate for Hebbian plasticity (*α*_*hebb*_) was set to 0.00005 for simulations in figure 4 and 0.0005 for simulations in figure extended data figure 6-1. Weights were normalized after Hebbian learning to ensure that the Euclidean norm of the incoming weights to each unit in layer two was equal to one. The model was “trained” over 100 trials in the same perceptual discrimination task described above and an additional 100 trials of the task were completed to measure emergent noise correlations in the hidden layer. Noise correlations were measured by regressing out variance attributable to the stimulus on each trial, and then computing the Pearson correlation of residual firing rate across each pair of neurons for the 100 testing trials (figure 4B&C).

### Learning readout in multiple discrimination task

In order to test the impact of contextual noise correlations on learning (Cohen and Newsome, 2008), the perceptual discrimination task was extended to include two dimensions and two interleaved trial types: one in which an up/down discrimination was performed (vertical), and one in which a right/left discrimination was performed (horizontal). Each trial contained motion on the vertical axis (up or down) and on the horizontal axis (left or right), but only one of these motion axes was relevant on each trial as indicated by a cue.

In order to model this task, we extended our two-layer feed-forward network to include 4 pools of input units, 4 output units, and 2 task units (figure 5A). Each homogenous pool of 100 input units encoded a conjunction of the movement directions (up-right, up-left, down-right, down-left). On each trial, the mean firing rate of each input unit population was determined according to their tuning preferences:

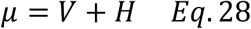

where V was +1/-1 for trials with the preferred/anti-preferred vertical motion direction H was +1/-1 for trials with the preferred/anti-preferred horizontal motion direction. Firing rates for individual neurons were sampled from a multivariate Gaussian distribution with mean *μ* and a covariance matrix that depended on trial type (vertical versus horizontal) and the level of same pool, relevant pool, and irrelevant pool correlations.

In order to create a covariance matrix, we stipulated a desired standard error of the mean for summed population activity (SEM=20 for simulations in figure 5) and determined the summed population variance that would correspond to that value 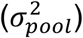. We then determined the variance on individual neurons that would yield this population response under a given noise correlation profile as follows:

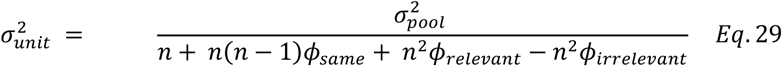

where *ϕ*_*same*_ is the level of same pool correlations (range: 0-0.2 in our simulations), *ϕ*_*relevant*_ is the level of relevant pool correlations (range: 0-0.2 in our simulations), *ϕ*_*relevant*_ is the level of irrelevant pool correlations (range: 0-0.2 in our simulations. Note that increasing the same pool or in pool correlations reduces the overall variance in order to preserve the same level of variance on the task relevant dimension in the population response, but that increasing irrelevant pool correlations has the opposite effect. Covariance elements of the covariance matrix were determined as follows:

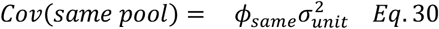

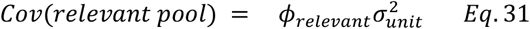

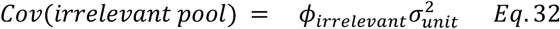

Variance and covariance values above were used to construct a covariance matrix for each trial type (vertical/horizontal) as depicted in figure 1.

Output units corresponded to the four possible task responses (up, down, left, right) and were activated according to a weighted sum of their inputs as described previously. Task units were modeled as containing perfect information about the task cue (vertical versus horizontal) and each task unit projected with strong fixed weights (1000) to both responses that were appropriate for that task. Decisions were made on each trial by selecting the output unit with the highest activity level. Weights to chosen output unit were updated using the same reinforcement learning procedure described in the two alternative perceptual learning task.

## Results

We examine how noise correlations affect learning in a simplified neural network where the appropriate readout of hundreds of weakly tuned units is learned over time through reinforcement. In order to isolate the effects of noise correlations on learning, rather than their effects on other factors such as representational capacity, we consider population encoding schemes at the input layer that can be constrained to a fixed signal-to-noise ratio. This assumption differs from previous work on noise correlations where the *variance* of the neural population is assumed to be fixed and covariance is changed to produce noise correlations, thereby affecting the representational capacity of the population (figure 2A; (Averbeck et al., 2006; Moreno-Bote et al., 2014)). Under our assumptions, a fixed signal-to-noise ratio can be achieved for any level of noise correlations by scaling the variance (figure 2B; equations 1-3), or, alternately scaling the magnitude of the signal (equation 25). While we do not discount the degree to which noise correlations affect the encoding potential of neural populations, we believe that in many cases the relevant information is limited by extrinsic factors (eg. the stimulus itself, or upstream neural populations providing input (Ecker et al., 2011; Beck et al., 2012; Kanitscheider et al., 2015)). Under such conditions, reducing noise correlations can increase information only until it saturates because all of the available incoming information is encoded. Beyond that, increasing encoding potential is not possible as it would be tantamount to the population “creating new information” that was not communicated by inputs to the population. Therefore, our framework can be thought of as testing how best to format limited available information in a neural population in order to ensure that an acceptable readout can be rapidly and robustly learned.

We propose that within this framework, noise correlations of the form that have previously been shown to limit encoding are beneficial because they constrain learning to occur over the most relevant dimensions. In general, a linear readout can be thought of as hyperplane serving as a classification boundary in an N dimensional space, where N reflects the number of neurons in a population. Learning in such a framework involves adjustments of the hyperplane to minimize classification errors. The most useful adjustments are in the dimension that best discriminates signal from noise (central arrows in figure 2C&D), but adjustments may also occur in dimensions orthogonal to the relevant one (such as “twisting” of the hyperplane depicted by curved arrows in figure 2C&D) that could potentially impair performance, or slow down learning. Our motivating hypothesis is that by focusing population activity into the task relevant dimension, noise correlations can increase the fraction of hyperplane adjustments that occur in the task relevant dimension (figure 2D), thus reducing the effective dimensionality of readout learning.

### Noise correlations enable faster learning in a fixed signal-to-noise regime

In order to test our overarching hypothesis, we constructed a fully connected two-layer feed-forward neural network in which input layer units responded to one of two stimulus categories (pool 1 & pool 2) and each output unit produced a response consistent with a category perception (left/right units in figure 3A). On each trial, the network was presented with one stimulus at random, and input firing for each pool was drawn from a multivariate Gaussian with a covariance that was manipulated while preserving the population signal-to-noise ratio. Output units were activated according to a weighted average of inputs and a response was selected according to output unit activations. On each trial, weights to the selected action were adjusted according to a reinforcement learning rule that strengthened connections that facilitated a rewarded action and weakened connections that facilitated an unrewarded action (Law and Gold, 2009).

Noise correlations led to faster and more robust learning of the appropriate stimulus-response mapping. All neural networks learned to perform the requisite discrimination, but neural networks that employed correlations among similarly tuned neurons learned more rapidly (figure 3B). After learning, networks that employed such noise correlations assigned more homogenous weights to input units of a given pool than did networks that lacked noise correlations (compare figure 3C&D). This led to better trained-task performance (figure 3E; Pearson correlation between noise correlations and test performance: R = 0.29, p < 10e-50) and greater robustness to adversarial noise profiles (figure 3F; R = 0.81, p < 10e-50) in the networks that employed noise correlations. Critically, these learning advantages emerged despite the fact that optimal readout of all networks achieved similar levels of performance and robustness (figure 3E&F, compare optimal readout across conditions).

### Learning benefits from noise correlations are greatest for large, low SNR populations

In order to better understand how noise correlations promoted faster learning we developed an analytical method for describing learning trajectories (see methods). Our method considered the impacts of two influences on weight updates over time: 1) weight updates in the signal dimension that tend to align with the signal and improve performance and 2) weight updates perpendicular to the signal dimension, which through chance alignment with trial-to-trial firing rate variability allow noise to impact decisions and therefore and hinder performance (fig 4a). Noise correlations implemented using our methods decreased the latter form of weight updates (fig 4b), leading updates in the signal dimension to more quickly dominate performance (fig 4c), thereby speeding analytical predictions for learning (fig 4d&e). The analytically derived learning advantage for fixed-SNR noise correlations was greatest for situations in which SNR was relatively low and neural populations were large (fig 4f).

The advantage of noise correlations for learning speed did not depend on specific assumptions about whether SNR was balanced by adjusting signal or noise. We employed an alternate method for creating fixed-SNR noise correlations that amplified signal, rather than reducing variance, in order to maintain SNR for higher levels of noise correlation (equation 25). Such noise correlations could be thought of as reflecting amplification of both signal and shared noise that would result from top down recurrent feedback (Haefner et al., 2016). Under such assumptions, noise correlations sped learning and led to more robust weight profiles, similarly to in our previous simulations (Extended data Fig 3-1).

### Noise correlations that do not maintain signal-to-noise ratio can introduce a tradeoff between learning speed and asymptotic performance

In contrast, our learning speed results depended critically on the assumption that signal-to-noise ratio is maintained across different levels of noise correlation. In order to test this dependency, we examined performance of a family of models that contained a single parameter allowing them to range in assumptions from fixed SNR (m=1) to fixed signal and single unit variance, analogous to assumptions of Averbeck and colleagues (Averbeck et al., 2006) (m=0). Consistent with our previous results, noise correlations improve learning in the m=1 case, and consistent with Averbeck 2006, asymptotic performance is reduced by noise correlations in the m=0 case (Fig 5). Interestingly, for intermediate assumptions between these two extremes, noise correlations promote faster learning improving performance in the short run, but at the cost of lower asymptotic accuracy. Thus, under such assumptions, adjusting noise correlations between similarly tuned neurons could potentially optimize a tradeoff between short-term gains from rapid learning and long term gains from higher asymptotic performance.

### Hebbian learning can produce useful noise correlation structure

Given that noise correlations implemented in our previous simulations, like those observed in the brain, depended on the tuning of individual units, we tested whether such noise correlations might be produced via Hebbian plasticity. Specifically, we considered an extension of our neural network in which an additional intermediate layer is included between input and output neurons (figure 6a). Input units were again divided into two pools that differed in their encoding, but variability was uncorrelated across neurons within a given pool. Connections between the input layer and intermediate layer were initialized such that each input unit strongly activated one intermediate layer unit, and shaped over time using a Hebbian learning rule that strengthened connections between co-activated neuron pairs. Despite the lack of noise correlations in the input layer of this network (figure 6b; mean[std] in pool residual correlation = 0.0015[0.10]), neurons in the intermediate layer developed tuning-specific noise correlations of the form that were beneficial for learning in the previous simulations (figure 6c; mean[std] in pool residual correlation = 0.55[0.07]; *t*-test on difference from input layer correlations *t* = 443, *dof* = 19800, *p* < 10e-50). Hebbian learning produced analogous noise correlation structure when initialized with random weights (Extended data figure 6-1). The ability of Hebbian learning to reduce the dimensionality of the input units is consistent with previous theoretical work showing that it extracts the first principal component of the input vector, which in this case, is the signal (Oja, 1982)

### Dynamic, task-dependent noise correlations speed learning by constraining it to relevant feature dimensions

In order to understand how noise correlations might impact learning in mixed encoding populations, we extended our perceptual discrimination task to include two directions of motion discrimination (eg. up/down and left/right). On each trial, a cue indicated which of two possible motion discriminations should be performed (figure 7A, left; (Cohen and Newsome, 2008)). We extended our neural network to include four populations of one hundred input units, each population encoding a conjunction of motion directions (up-right, up-left, down-right, down-left; figure 7A; input layer). Two additional inputs provided a perfectly reliable “cue” regarding the relevant feature for the trial (figure 7A; task units). Four output neurons encoded the four possible responses (up, left, down, right) and were fully connected to the input layer (figure 7A; output layer). Task units were hard wired to eliminate irrelevant task responses, but weights of input units were learned over time as in our previous simulations.

**Figure 7:**
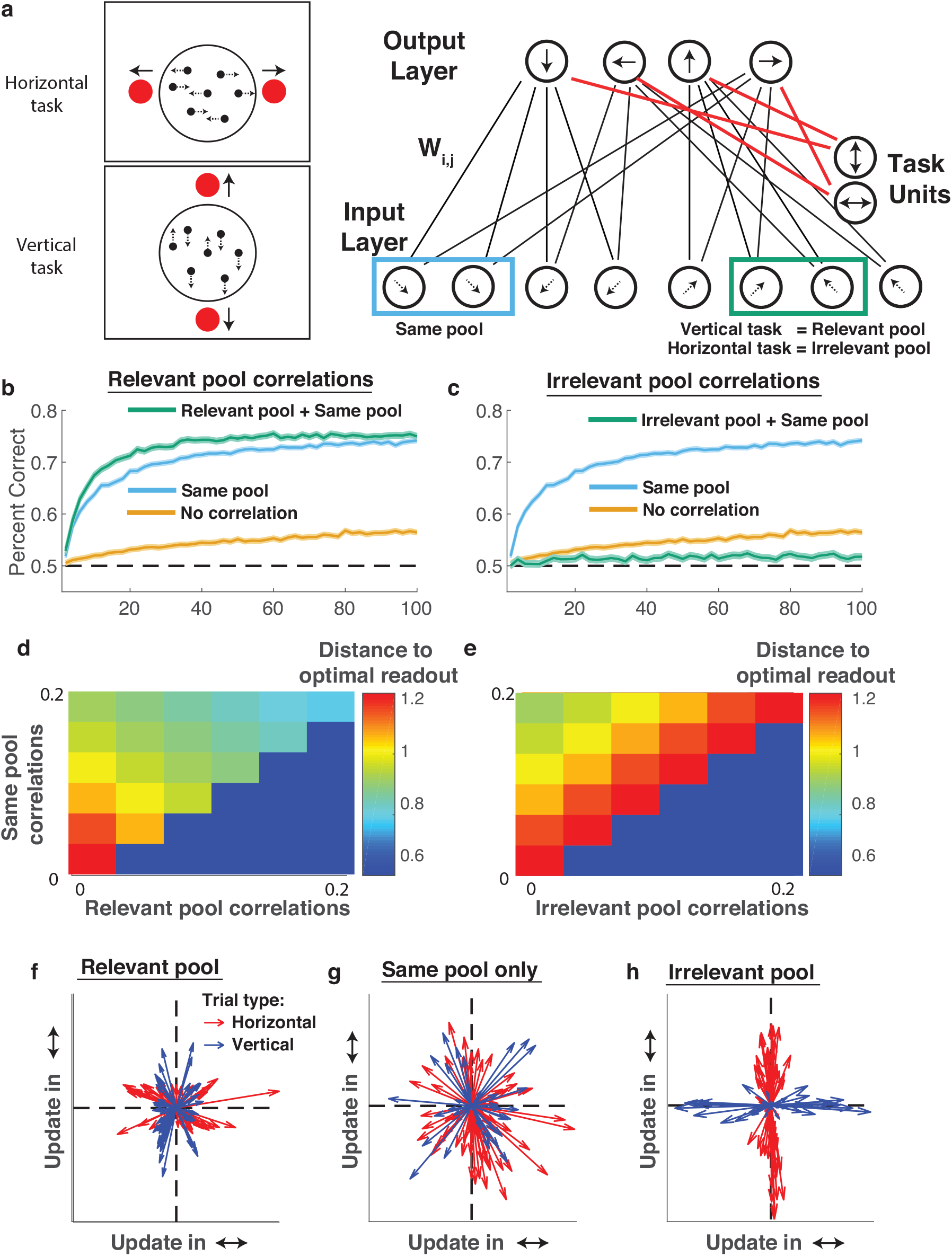
Task dependent noise correlations affect learning speed by projecting learning onto specific feature dimensions. **A)** A neural network was trained to perform two interleaved motion discrimination tasks (left; (Cohen and Newsome, 2008)). Network schematic (right) depicts two-layer feed-forward network in which each homogenous pool of input units represents two dimensions of motion (up versus down, and left versus right), and output units produce responses in favor of alternative actions (up, down, left, right). Each homogenous pool of input units is identically tuned to one of four conjunctions of movement directions: up-left, down-left, up-right, down-right. Two additional input units provide cue information that biases output units to produce an output corresponding to the discrimination appropriate on this trial (eg. horizontal or vertical). Noise correlations were manipulated among 1) identically tuned neurons (blue rectangle; same pool), 2) neurons that have similar encoding of the task relevant feature (green rectangle pair in vertical trials; relevant pool), and 3) neurons that have similar encoding of the task irrelevant feature (green rectangle pair in horizontal trials; irrelevant pool). **B&C**) Learning curves showing accuracy (ordinate) over trials (abscissa) for models 1) lacking noise correlations (orange), 2) containing noise correlations that are limited to neurons that have same tuning for both features (same pool; blue), 3) containing same pool noise correlations along with correlations between neurons in different pools that have the same tuning for the task-relevant feature (in pool+rel pool; green in **B**), and 4) containing in-pool noise correlations along with correlations between neurons in different pools that have the same tuning for the task irrelevant feature (in pool+irrel pool; green in **C**). **D&E**) Distance between learned weights and the optimal readout (color) for models that differ in their level of “in pool” correlations (ordinate, both plots), “relevant pool” correlations (abscissa, **D**), and “irrelevant pool” correlations (abscissa, **E**). **F**,**G**,**H**) Weight updates for example learning sessions were projected into a two dimensional space in which net updates to the relative contribution of vertical motion information (eg. up versus down) is represented on the abscissa and updates to the relative contribution of horizontal motion information (eg. left versus right) is represented on the ordinate. Arrows reflect single trial weight updates and are colored according to the trial type (red = horizontal discrimination, blue = vertical discrimination). Weight updates for a model with only “in pool” correlations look similar across trial types (**G**), but weight updates for a model with “relevant pool” correlations indicate more weight updating on the relevant feature (**F**), whereas the opposite was observed in the case of “irrelevant pool” correlations (**H**).

Learning performance in the two-feature discrimination task depended not only on the level of noise correlations, but also on the type. As in the previous simulation, adding noise correlations to each individual population of identically tuned units led to faster learning of the appropriate readout (Figure 7B&C, compare blue and yellow; Figure 7D&E, vertical axis; mean[std] accuracy across training: 0.54[0.05] and 0.70[0.05] for minimum (0) and maximum (0.2) in pool correlations, t-test for difference in accuracy: *t* = 226, *dof* = 19998, *p* <10e-50).

However, the more complex task design also allowed us to test whether dynamic trial-to-trial correlations might further facilitate learning. Specifically, correlations that increase shared variability among units that contribute evidence to the same response have been observed previously (Cohen and Newsome, 2008), and could in principle focus learning on relevant dimensions (figure 2C&D) even when those dimensions change from trial to trial. Indeed, adding correlations among separate pools that share the same encoding of the relevant feature (eg. UP on a vertical trial) led to faster learning (figure 7B; mean[std] training accuracy for model with relevant pool correlations: 0.73[0.05], *t*-test for difference from in pool correlation only model: *t* = 34, *dof* =19998, *p* <10e-50) and weights that more closely approached the optimal readout (figure 7E, horizontal axis). In contrast, when positive noise correlations were introduced across separate encoding pools that shared the same tuning for the irrelevant dimension on each trial (eg. UP on a horizontal trial) learning was impaired dramatically (figure 7C; mean[std] training accuracy for model with irrelevant pool correlations: 0.51[0.05], *t*-test for difference from in pool correlation only model: *t* = −278, *dof* =19998, *p* <10e-50) and learned weights diverged from the optimal readout (figure 7F, horizontal axis). Model performance differences were completely attributable to learning the readout, as all models performed similarly when using the optimal readout (extended data figure 7-1).

In order to test the idea that noise correlations might focus learning onto relevant dimensions, we extracted weight updates from each trial and projected these updates into a two-dimensional space where the first dimension captured the relative sensitivity to leftward versus rightward motion and the second dimension captured relative sensitivity to upward versus downward motion. In the model where input units were only correlated within their identically tuned pool, weight updates projected in all directions more or less uniformly (figure 7G), and did not differ systematically across trial types (vertical versus horizontal). However, dynamic noise correlations that shared variability across the relevant dimension tended to push weight updates onto the appropriate dimension for a given trial (figure 4F; *t*-test for difference in the magnitude of updating in up/down and left/right dimensions across conditions [up/down – left/right]: *t* = 3.4, *dof*=98, *p* = 0.001). In contrast, dynamic noise correlations that shared variability across the irrelevant dimension tended to push weight updates onto the wrong dimension (figure 4H; t-test for difference in the magnitude of updating in up/down and left/right dimensions across conditions [up/down – left/right]: *t* = −9.5, *dof*=98, *p* = 10e-14). Both of these trends were consistent across simulations, providing an explanation for the performance improvements achieved by relevant noise correlations (projection of learning onto an appropriate dimension) and performance impairments produced by irrelevant noise correlations (projection of learning onto an inappropriate dimension).

## Discussion

Taken together, our results suggest that in settings where the population signal-to-noise ratio is limited by external factors (eg. inputs) and relevant task representations are low dimensional, noise correlations can make learning faster and more robust by focusing learning on the most relevant dimensions. We demonstrate this basic principle in a simple perceptual learning task (figure 3), where beneficial noise correlations between similarly tuned units could be produced through a simple Hebbian learning rule (figure 6). We extended our framework to a contextual learning task to demonstrate that dynamic noise correlations that bind task relevant feature representations facilitate faster learning (figure 7b&d) by pushing learning onto task-relevant dimensions (figure 7f). Given the pervasiveness of noise correlations among similarly tuned sensory neurons (Zohary et al., 1994; Maynard et al., 1999; Bair et al., 2001; Averbeck and Lee, 2003; Cohen and Maunsell, 2009; Huang and Lisberger, 2009; Ecker et al., 2010; Gu et al., 2011; Adibi et al., 2013), and that the noise correlations dynamics beneficial for learning in our simulations are similar to those that have been observed *in vivo* (Cohen and Newsome, 2008), we interpret our results as suggesting that noise correlations between similarly tuned neurons are a feature of neural coding architectures that ensures efficient readout learning, rather than a bug that limits encoding potential.

This interpretation rests on several assumptions in our model. Of particular importance is the assumption that signal-to-noise ratio of our populations is fixed, meaning that our manipulation of noise correlations can focus variance on specific dimensions without gaining or losing information. This assumption reflects conditions in which information is limited at the level of the inputs to the population, for instance due to noisy peripheral sensors (Beck et al., 2012; Kanitscheider et al., 2015). In such conditions, even with optimal encoding, population information saturates at an upper bound determined by the information available in the inputs to the population. Therefore, fixing the signal-to-noise ratio enabled us to examine the effect of noise correlations on downstream processes that learn to read-out the population code in the absence of any influence of noise correlations on the quantity of information contained within that population code.

Previous theoretical work exploring the role of noise correlations in encoding has typically assumed that single neurons have a fixed variance, such that tilting the covariance of neural populations towards or away from the dimension of signal encoding would have a large impact on the amount of information that can be encoded by a population (figure 1a; (Averbeck et al., 2006; Moreno-Bote et al., 2014)). Such assumptions lead to the idea that positive noise correlations among similarly tuned neurons limit encoding potential, raising the question of why they are so common in the brain (Cohen and Kohn, 2011). In considering the implications of this framework, one important question is: if information encoded by the population can be increased by changing the correlation structure among neurons, where does this additional information come from? In some cases, the neural population in question may indeed receive sufficient task relevant information from upstream brain regions to reorganize its encoding in this way, but in other cases it is likely that information is limited by the inputs to a neural population (Kanitscheider et al., 2015; Kohn et al., 2016). In cases where incoming information is limited, further increasing representational capacity is not possible, and formatting information for efficient readout is essentially the best that the population code could do. Here we show that the noise correlations that have previously been described as “information limiting” are exactly the type of correlations that format information most efficiently for readout learning under such conditions.

Between these two bookends of a fixed signal-to-noise ratio and fixed single unit variance, we also simulated intermediate regimes which do not perfectly preserve the signal-to-noise ratio. In these intermediate regimes, a tradeoff emerges: noise correlations between similarly tuned neurons produce faster learning in the short term, at the cost of lower levels of asymptotic performance in the long run (Fig 5).

Jointly considering these perspectives on noise correlations provides a more nuanced view of how neural representations are likely optimized for learning. In order to optimize an objective function, a neural population can reduce correlated noise in task relevant dimensions to increase its representational capacity up to some level constrained by its inputs (Figure 8, left). But once the population is fully representing all task relevant information that has been provided to it, it can additionally optimize representations by pushing as much variance onto task relevant dimensions as possible, thereby affording efficient learning in downstream neural populations (Figure 8, right). In short, optimization of a neural population code does not occur in a vacuum, and instead depends critically on both upstream (eg. input constraints) and downstream (eg. readout) neural populations (Figure 8). In this view, if a neural population is *not* fully representing the decision relevant information made available to it, then learning could improve the efficiency of representations by reducing rate limiting noise correlations as has been observed in some paradigms (Gu et al., 2011; Ni et al., 2018). In contrast, once available information is fully represented, readout learning could be further optimized by reformatting population codes such that variability is shared across neurons with similar tuning for the relevant task feature, producing the sorts of dynamic noise correlations that have been observed in well trained animals (Cohen and Newsome, 2008).

**Figure 8:**
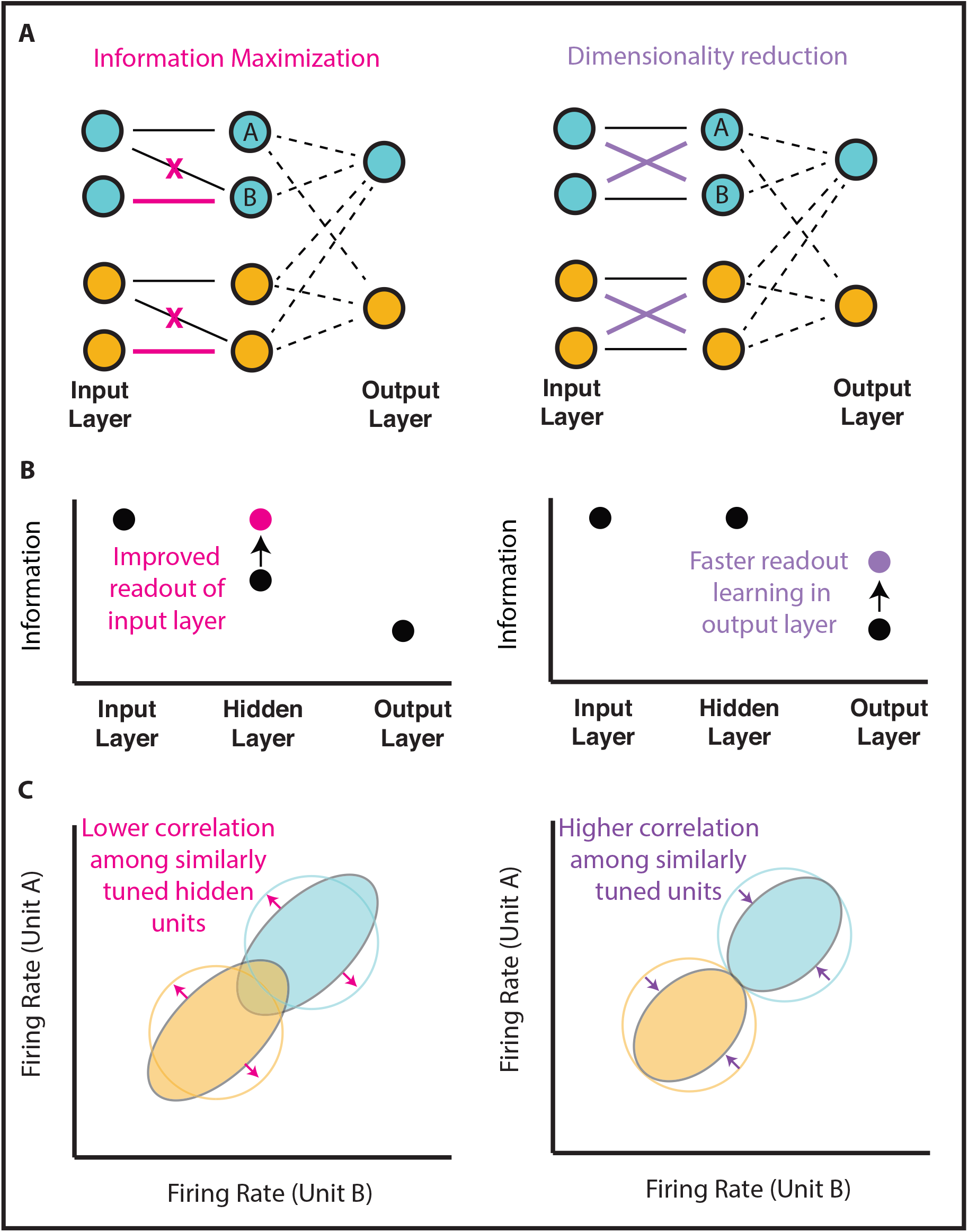
Information maximization and dimensionality reduction can be useful for learning under different situations and have opposite effects on noise correlations among similarly tuned units. **A)** A schematic representation of a three layer neural network in which units provide evidence for one of two categorizations (blue/orange). In the left network, the hidden layer initially has access to information from only one of two independent units in each pool, but weights are subsequently adjusted to increase task-relevant information represented in the hidden layer (pink). In the right network, the hidden layer initially has access to all task-relevant information, but weights are subsequently adjusted to share signal and noise across similarly tuned units to afford dimensionality reduction (purple). Note that the information maximizing weight adjustments (left, pink) increase signal-to-noise ratio in the hidden layer but preserve the variance in firing rate of individual neurons, whereas the dimensionality reducing weight adjustments (right, purple) maintain a fixed signal-to-noise ratio in hidden units, but decrease the variance of individual units by averaging across multiple similarly tuned inputs. Dashed lines to output units reflect weights that need to be learned based on feedback. **B)** Task relevant information (mutual information between unit activations and stimulus category; abscissa) is depicted for each layer (ordinate). Weight adjustments affording information maximization (left) increase task relevant information in the hidden layer (pink), whereas weight adjustments that afford dimensionality reduction (right) do not affect task-relevant information in the hidden layer itself but instead increase the rate of learning in the output layer, thereby leading to more task-relevant information in the output layer (purple). **C)** Weight adjustments for information maximization (pink in panel A) *decrease* correlations among hidden units A&B by removing shared input from a single input unit and instead providing independent sources of input to each unit (pink arrows). In contrast, weight adjustments for dimensionality reduction *increase* noise correlations among hidden units A&B by providing them with the same mixture of information from the two identically tuned input units. We propose that both of these processes play a critical role in learning and that changes in noise correlations across learning will depend critically on which process dominates. As shown in panel B, this will depend critically on whether the neural population in question has already fully represented information available from its inputs. In principle, these processes could occur serially, with early learning maximizing information available in intermediate layers (left) and later learning compressing that information into a format allowing rapid readout learning (right).

In addition to key assumptions about an external limitation on signal-to-noise, our modeling included a number of simplifying assumptions that are unlikely to hold up in real neural populations. For example, we consider pools of neurons identically tuned to discrete stimuli, rather than a continuous space of stimuli and the heterogeneous populations observed in sensory cortical regions of the brain. Previous work has shown that noise correlations do not necessarily limit encoding potential in heterogenous populations with diverse tuning (Shamir and Sompolinsky, 2004; 2006; Chelaru and Dragoi, 2008; Ecker et al., 2011). A primary goal of our work was to identify the computational principles that control the speed at which readout can be learned, and our simplified populations are considerably more tractable and transparent than realistic neural populations. The principles that we identify here are certainly at play in real neural populations, albeit with implications that are far less transparent. In particular, in a population with diverse tuning profiles, the degree to which individual neurons are informative about a task-relevant discrimination will vary. To benefit learning through coordinated weighted changes, the correlation structure in such populations should reflect this variability. Empirical studies in macaques suggest that such variability is indeed present in real populations (Cohen and Maunsell, 2009; Rabinowitz et al., 2015; Ruff and Cohen, 2016; Bondy et al., 2018). We hope that our simplified results pave the way for future work to assess nuances that can emerge in mixed heterogeneous populations, or in more realistic architectures that go beyond the simple feed forward flow of information considered here.

### Model predictions

Our work shows that noise correlations can focus the gradient of learning onto the most appropriate dimensions. Thus, our model predicts that the degree to which similarly tuned neurons are correlated during a perceptual discrimination should be positively related to performance improvements experienced on subsequent discriminations. In contrast, our model predicts that the degree of correlation between neurons that are similarly tuned to a task irrelevant feature should control the degree of learning on irrelevant dimensions, and thus negatively relate to performance improvements on subsequent discriminations. These predictions are strongest for the earliest stages of learning where weight adjustments are critical for subsequent performance, but they may also hold for later stages of learning, when correlations on irrelevant dimensions, including independent noise channels, could potentially lead to systematic deviations from optimal readout (figure 2f, 4d&e). These predictions could be tested by recording neural responses to a stimulus set that differs across multiple features to characterize both signal-to-noise and correlated variability for each feature discrimination. A strong prediction of our model is that correlated variability within neurons tuned to a given feature should be a predictor of subsequent learning of responses to that feature – above and beyond feature value discriminability.

One interesting special case involves tasks where the relevant dimension changes in an unsignaled manner (Birrell and Brown, 2000). In such tasks, noise correlations on the previously relevant dimension would, after such an “extradimensional shift”, force gradients into a task-irrelevant dimension and thus impair learning performance. Interestingly, learning after extra-dimensional shifts can be selectively improved by enhancing noradrenergic signaling (Devauges and Sara, 1990; Lapiz and Morilak, 2006), which leads to increased arousal (Joshi et al., 2016; Reimer et al., 2016) and decreased cortical pairwise noise correlations in sensory and higher order cortex (Vinck et al., 2015; Joshi and Gold, n.d.). While these observations have been made in different paradigms, our model suggests that the reduction of noise correlations resulting from increased sustained levels of norepinephrine after an extradimensional shift (Bouret and Sara, 2005) could mediate faster learning by expanding the dimensionality of the learning gradients (compare figure 7G to 7F) to consider features that have not been task-relevant in the past.

### Relation to attentional effects on noise correlations

In broad strokes, our finding that manipulation of noise correlations can focus variance on specific dimensions is in line with specific models of attention. In particular, noise reduction in task irrelevant dimensions might be considered in the same light that is often cast on suppression of task irrelevant dimensions by attentional mechanisms (Zanto and Gazzaley, 2009), in particular for purposes of accurate credit assignment (Akaishi et al., 2016; Leong et al., 2017). One possibility is that compressed low-dimensional task representations in higher-order decision regions (Mack et al., 2019) may pass accumulated decision related information back to sensory regions in order to approximate Bayesian inference (Haefner et al., 2016; Bondy et al., 2018; Lange et al., 2018). As task relevant features are learned, such a process would promote noise correlations between neurons coding those relevant features. In other words, noise correlations may reflect a chosen hypothesis about which feature is relevant for predicting outcomes. Such a signal would be beneficial if it could persist (and thus preserve correlations between neurons tuned to the same task relevant feature value) until the time of feedback or reinforcement. Recent work showing strengthened noise correlations between similarly tuned neurons during working memory maintenance suggests that this might very well be the case (Merrikhi et al., 2018).

One observation that seems at odds with this interpretation is that manipulations of attention that cue a particular location or feature tend to decrease noise correlations among neurons that encode that location or feature (Cohen and Maunsell, 2009; Mitchell et al., 2009; Cohen and Maunsell, 2011; Herrero et al., 2013; Doiron et al., 2016). The effects of attentional cuing on noise correlations are dynamic in that cues change from one trial to the next, and contextual, in that noise correlations are reduced most dramatically among neurons that contribute evidence toward the same response in a manner consistent with increasing the amount of task relevant information in the population code (Ruff and Cohen, 2014; Downer et al., 2015). These effects are generally observed in well-trained animals during task performance and may not result from the same processes as the longer timescale noise correlation structure. Indeed, there may be a tradeoff between learning and performance, particularly if the computations giving rise to noise correlations do so without perfectly preserving signal-to-noise ratio. Our model does not account for these attentional effects, as we intentionally constrained the signal-to-noise ratio of our neural populations, thereby eliminating any potential changes in information encoding potential. However, we hope that our work motivates future studies to jointly consider the impacts of noise correlations on both learning and immediate performance in order to better understand the potentially competing imperatives that the brain faces in dynamically controlling the correlation structure of its own representations (see (Haimerl et al., 2019) for one attempt to do so).

### Origins of useful noise correlations

One important question stemming from our work is how noise correlations emerge in the brain. This question has been one of longstanding debate, largely because there are so many potential mechanisms through which correlations could emerge (Kanitscheider et al., 2015; Kohn et al., 2016). Noise correlations could emerge from convergent and divergent feed forward wiring (Shadlen and Newsome, 1998), local connectivity patterns within a neural population (Hansen et al., 2012; Smith et al., 2013), or top down inputs provided separately to different neural populations (Haefner et al., 2016). Here we show that static noise correlations that are useful for perceptual learning emerge naturally from Hebbian learning in a feed-forward network. While this certainly suggests that useful noise correlations could emerge through feed forward wiring, it is also possible to consider our Hebbian learning as occurring in a one-step recurrence of the input units, and thus the same data support the possibility of noise correlations through local recurrence. The context dependent noise correlations that speed learning (figure 7), however, would not arise through simple Hebbian learning. Such correlations could potentially be produced through selective top-down signals from the choice neurons, as has been previously proposed (Wimmer et al., 2015; Haefner et al., 2016; Bondy et al., 2018; Lange et al., 2018). Moreover, top-down input may selectively target neuronal ensembles produced through Hebbian learning (Collins and Frank, 2013). While previous work has suggested that such a mechanism could be adaptive for accumulating information over the course of a decision (Haefner et al., 2016), our work demonstrates that the same mechanism could effectively be used to tag relevant neurons for weight updating between trials, making efficient use of top-down circuitry. Haimerl et al. recently made a similar point, showing that stochastic modulatory signals shared across task-informative neurons can serve to tag them for a decoder (Haimerl et al., 2019).

### Noise correlations as inductive biases

Artificial intelligence has undergone a revolution over the past decade leading to human level performance in a wide range of tasks (Mnih et al., 2015). However, a major issue for modern artificial intelligence systems, which build heavily on neural network architectures, is that they require far more training examples than a biological system would (Hassabis et al., 2017). This biological advantage occurs despite the fact that the total number of synapses in the human brain, which could be thought of as the free parameters in our learning architecture, is much greater than the number of weights in even the most parameter-heavy deep learning architectures. Our work provides some insight into why this occurs; correlated variability across neurons in the brain constrain learning to specific dimensions, thereby limiting the effective complexity of the learning problem (figures 4A, 7F-G). We show that, for simple tasks, this can be achieved using Hebbian learning rules (figure 6), but that contextual noise correlations, of the form that might be produced through top-down signals (Haefner et al., 2016), are critical for appropriately focusing learning in more complex circumstances. In principle, algorithms that effectively learn and implement noise correlations might reduce the amount of data needed to train AI systems by limiting degrees of freedom to those dimensions that are most relevant. Furthermore, our work suggests that large scale neural recordings in early stages of learning complex tasks might serve as indicators of the inductive biases that constrain learning in biological systems.

In summary, we show that under external constraints of task-relevant information, noise correlations that have previously been called “rate limiting” can serve an important role in constraining learning to task-relevant dimensions. In the context of previous theory focusing on representation, our work suggests that neural populations are subject to competing forces when optimizing covariance structures; on one hand reducing correlations between pairs of similarly tuned neurons can be helpful to fully represent available information, but increasing correlations among similarly tuned neurons can be helpful for assigning credit to task relevant features. We believe that this view of the learning process not only provides insight to understanding the role of noise correlations in the brain, but opens up the door to better understand the inductive biases that guide learning in biological systems.

## Acknowledgements

We would like to thank Josh Gold, Rex Liu, Michael Frank, Drew Linsley, Chris Moore and Jan Drugowitsch for helpful discussion. This work was funded by NIH grants F32MH102009 and R00AG054732 (MRN), NINDS R21NS108380 (AB). The funders had no role in study design, data collection and analysis, decision to publish or preparation of the manuscript.

## Extended data

**Extended data figure 3-1:**
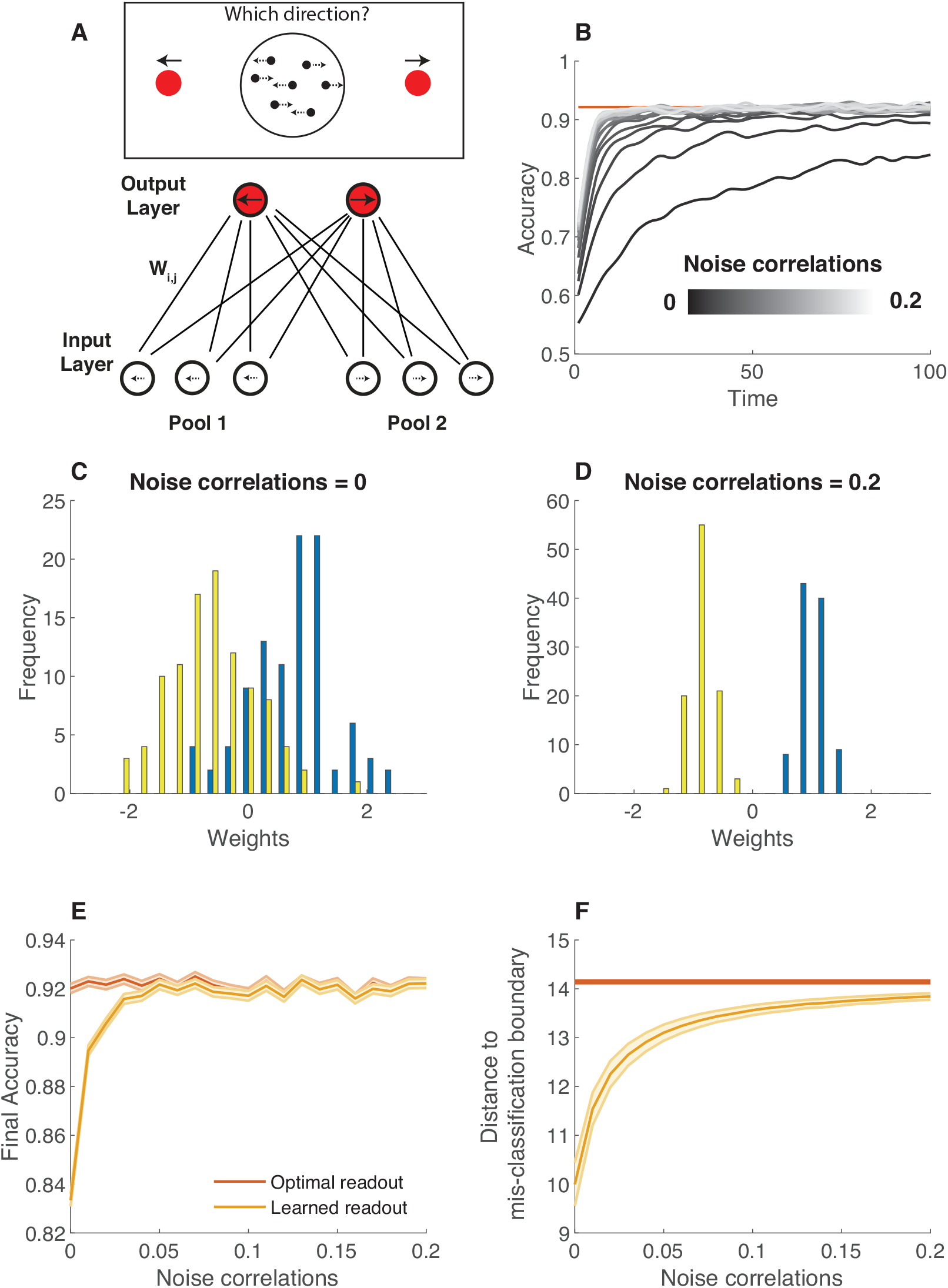
Noise correlations that maintain signal-to-noise ratio by scaling signal lead to faster and more robust learning of a perceptual discrimination. This figure is a replication of results reported in figure 3 of the main text, except that noise correlations are produced using equation 25 such that each unit maintains the same fixed variance across noise correlation conditions, and signal is scaled to maintain a fixed signal-to-noise ratio. **A)** A two-layer feed-forward neural network was designed to solve a two alternative forced choice motion discrimination task at or near perceptual threshold. Input layer contains two pools of neurons that provide evidence for alternate percepts (eg. leftward motion versus rightward motion) and output neurons encode alternate courses of actions (eg. saccade left versus saccade right). Layers are fully connected with weights randomized to small values and adjusted after each trial according to rewards (see methods). **B)** Average learning curves for neural network models in which population signal-to-noise ratio in pools 1 and 2 were fixed, but noise correlations (grayscale) were allowed to vary from small (dark) to large (light) values. **C&D)** Weight differences (left output – right output) for each input unit (color coded according to pool) after 100 timesteps of learning for low (**C**) and high (**D**) noise correlations. **E**) Accuracy in the last 20 training trials is plotted as a function of noise correlations for learned readouts (orange) and optimal readout (red). Lines/shading reflect Mean/SEM. F) The shortest distance, in terms of neural activation, required to take the mean input for a given category (eg. left or right) to the boundary that would result in misclassification is plotted for the final learned (orange) and optimal (red) weights for each noise correlation condition (abscissa). Lines/shading reflect Mean/SEM.

**Extended data figure 6-1:**
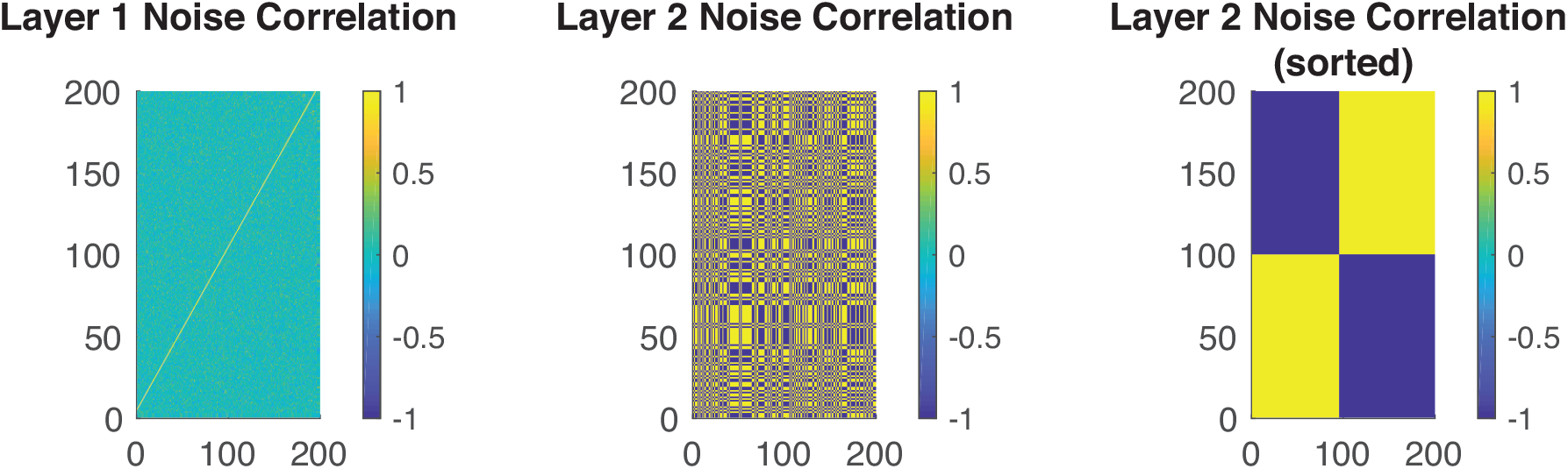
Emergence of noise correlations from Hebbian learning does not depend on weight initialization. In order to test whether beneficial noise correlations might have emerged in our Hebbian learning simulations due to our initialization biasing one-to-one connectivity between the input and hidden layers (see figure 4), we repeated these simulations with in a network that was initialized with random normal weight projects from layer 1 to layer 2. Simulations included performance of 200 trials with noise correlations measured in the final 100 trials of the simulation. Activity of layer 1 units was defined by a multivariate Gaussian with zero covariance elements, and thus it is not surprising that pairwise noise correlations measured in the activity of that layer were near zero (Left). Activity of layer 2 units was sculpted through Hebbian learning that shaped connectivity between layer 1 and layer 2. This learning led many pairs of neurons in layer 2 to become highly correlated and many other pairs to become anti-correlated (middle, blue and yellow elements, respectively). In order to understand the structure defining these correlations, we sorted the layer 2 units according to their relative projections to the two output units (weight to output 1 – weight to output 2), and recomputed the pairwise correlations (right). This sorting reveals that layer 2 units projecting to the same output unit are positively correlated with one another, whereas they negatively correlate with the layer 2 units that project to the opposing output neuron (note block diagonal structure).

**Extended data figure 7-1:**
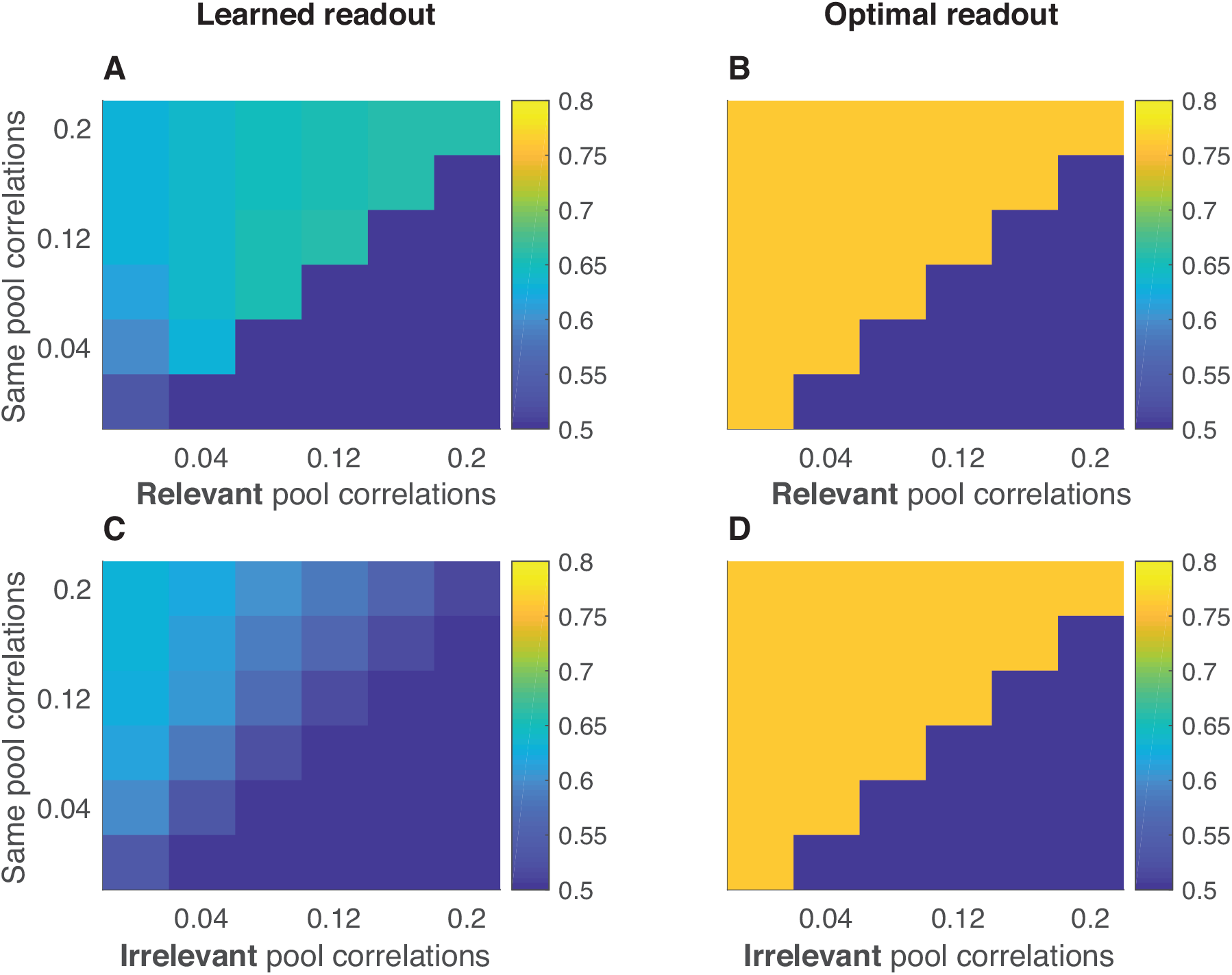
Noise correlations affect speed of learning, but not performance using optimal readout in multiple discrimination task. **A)** Mean test accuracy (color) of all models spanning the range of in pool correlations (abscissa) and relevant pool correlations (ordinate). **B)** Mean accuracy of same models using optimal readout, rather than the learned readout. **C)** Mean test accuracy (color) of all models spanning the range of in pool correlations (abscissa) and irrelevant pool correlations (ordinate). **D)** Mean accuracy of same models using optimal readout, rather than the learned readout. Note that performance of all models is identical when readout is optimal, rather than learned.

## References

Adibi M, McDonald JS, Clifford CWG, Arabzadeh E (2013) Adaptation improves neural coding efficiency despite increasing correlations in variability. Journal of Neuroscience 33:2108–2120.

Akaishi R, Kolling N, Brown JW, Rushworth M (2016) Neural Mechanisms of Credit Assignment in a Multicue Environment. Journal of Neuroscience 36:1096–1112.

Averbeck BB, Latham PE, Pouget A (2006) Neural correlations, population coding and computation. Nature Reviews Neuroscience 7:358–366.

Averbeck BB, Lee D (2003) Neural noise and movement-related codes in the macaque supplementary motor area. Journal of Neuroscience 23:7630–7641.

Bair W, Zohary E, Newsome WT (2001) Correlated firing in macaque visual area MT: time scales and relationship to behavior. Journal of Neuroscience 21:1676–1697.

Beck JM, Ma WJ, Pitkow X, Latham PE, Pouget A (2012) Perspective. Neuron 74:30–39.

Birrell JM, Brown VJ (2000) Medial frontal cortex mediates perceptual attentional set shifting in the rat. Journal of Neuroscience 20:4320–4324.

Bondy AG, Haefner RM, Cumming BG (2018) Feedback determines the structure of correlated variability in primary visual cortex. Nature Publishing Group:1–15.

Bouret S, Sara SJ (2005) Network reset: a simplified overarching theory of locus coeruleus noradrenaline function. Trends in Neurosciences 28:574–582.

Chelaru MI, Dragoi V (2008) Efficient coding in heterogeneous neuronal populations. Proceedings of the National Academy of Sciences 105:16344–16349.

Cohen MR, Kohn A (2011) Measuring and interpreting neuronal correlations. Nature Publishing Group 14:811–819.

Cohen MR, Maunsell JHR (2009) Attention improves performance primarily by reducing interneuronal correlations. Nature Publishing Group 12:1594–1600.

Cohen MR, Maunsell JHR (2011) Using neuronal populations to study the mechanisms underlying spatial and feature attention. Neuron 70:1192–1204.

Cohen MR, Newsome WT (2008) Context-Dependent Changes in Functional Circuitry in Visual Area MT. Neuron 60:162–173.

Collins AGE, Frank MJ (2013) Cognitive control over learning: creating, clustering, and generalizing task-set structure. Psychological Review 120:190–229.

Devauges V, Sara SJ (1990) Activation of the noradrenergic system facilitates an attentional shift in the rat. Behavioural Brain Research 39:19–28.

Doiron B, Litwin-Kumar A, Rosenbaum R, Ocker GK, Josić K (2016) The mechanics of state-dependent neural correlations. Nature Publishing Group 19:383–393.

Downer JD, Niwa M, Sutter ML (2015) Task engagement selectively modulates neural correlations in primary auditory cortex. Journal of Neuroscience 35:7565–7574.

Ecker AS, Berens P, Keliris GA, Bethge M, Logothetis NK, Tolias AS (2010) Decorrelated neuronal firing in cortical microcircuits. Science 327:584–587.

Ecker AS, Berens P, Tolias AS, Bethge M (2011) The effect of noise correlations in populations of diversely tuned neurons. Journal of Neuroscience 31:14272–14283.

Gu Y, Liu S, Fetsch CR, Yang Y, Fok S, Sunkara A, DeAngelis GC, Angelaki DE (2011) Perceptual learning reduces interneuronal correlations in macaque visual cortex. Neuron 71:750–761.

Haefner RM, Pietro Berkes, Fiser J (2016) Perceptual Decision-Making as Probabilistic Inference by Neural Sampling. Neuron 90:649–660.

Haimerl C, Savin C, Simoncelli EP (2019) Flexible and accurate decoding of neural populations through stochastic comodulation. Biorxiv 21:598.

Hansen BJ, Chelaru MI, Dragoi V (2012) Correlated variability in laminar cortical circuits. Neuron 76:590–602.

Hassabis D, Kumaran D, Summerfield C, Botvinick M (2017) Neuroscience-Inspired Artificial Intelligence. Neuron 95:245–258.

Hawkey DJC, Amitay S, Moore DR (2004) Early and rapid perceptual learning. Nature Publishing Group 7:1055–1056.

Herrero JL, Gieselmann MA, Sanayei M, Thiele A (2013) Attention-induced variance and noise correlation reduction in macaque V1 is mediated by NMDA receptors. Neuron 78:729–739.

Hu, Y., Zylberberg, J., & Shea-Brown, E. (2014). The sign rule and beyond: boundary effects, flexibility, and noise correlations in neural population codes. PLoS Comput Biol, 10(2), e1003469.

Huang X, Lisberger SG (2009) Noise correlations in cortical area MT and their potential impact on trial-by-trial variation in the direction and speed of smooth-pursuit eye movements. Journal of Neurophysiology 101:3012–3030.

Joshi S, Gold JI (n.d.) Context-Dependent Relationships between Locus Coeruleus Firing Patterns and Coordinated Neural Activity in the Anterior Cingulate Cortex. Biorxiv.

Joshi S, Li Y, Kalwani RM, Gold JI (2016) Relationships between Pupil Diameter and Neuronal Activity in the Locus Coeruleus, Colliculi, and Cingulate Cortex. Neuron 89:221–234.

Kanitscheider I, Coen-Cagli R, Pouget A (2015) Origin of information-limiting noise correlations. Proceedings of the National Academy of Sciences 112:E6973–E6982.

Kohn A, Coen-Cagli R, Kanitscheider I, Pouget A (2016) Correlations and Neuronal Population Information. Annu Rev Neurosci 39:237–256.

Krotov D, Hopfield JJ (2019) Unsupervised learning by competing hidden units. Proceedings of the National Academy of Sciences 116:7723–7731.

Lange RD, Chattoraj A, Beck JM, Yates JL, Haefner RM (2018) A confirmation bias in perceptual decision-making due to hierarchical approximate inference. Biorxiv.

Lapiz MDS, Morilak DA (2006) Noradrenergic modulation of cognitive function in rat medial prefrontal cortex as measured by attentional set shifting capability. Neuroscience 137:1039–1049.

Law C-T, Gold JI (2009) Reinforcement learning can account for associative and perceptual learning on a visual-decision task. Nature Neuroscience 12:655–663.

Leong YC, Radulescu A, Daniel R, DeWoskin V, Niv Y (2017) Dynamic Interaction between Reinforcement Learning and Attention in Multidimensional Environments. Neuron 93:451–463.

Mack ML, Preston AR, Love BC (2019) Ventromedial prefrontal cortex compression during concept learning. Nature Communications:1–11.

Maynard EM, Hatsopoulos NG, Ojakangas CL, Acuna BD, Sanes JN, Normann RA, Donoghue JP (1999) Neuronal interactions improve cortical population coding of movement direction. Journal of Neuroscience 19:8083–8093.

Merrikhi Y, Clark K, Noudoost B (2018) Concurrent influence of top-down and bottom-up inputs on correlated activity of Macaque extrastriate neurons. Nature Communications 9:5393.

Mitchell JF, Sundberg KA, Reynolds JH (2009) Spatial attention decorrelates intrinsic activity fluctuations in macaque area V4. Neuron 63:879–888.

Mnih V, Kavukcuoglu K, Silver D, Rusu AA, Veness J, Bellemare MG, Graves A, Riedmiller M, Fidjeland AK, Ostrovski G, Petersen S, Beattie C, Sadik A, Antonoglou I, King H, Kumaran D, Wierstra D, Legg S, Hassabis D (2015) Human-level control through deep reinforcement learning. Nature 518:529–533.

Moreno-Bote R, Beck J, Kanitscheider I, Pitkow X, Latham P, Pouget A (2014) Information-limiting correlations. Nature Publishing Group 17:1410–1417.

Ni AM, Ruff DA, Alberts JJ, Symmonds J, Cohen MR (2018) Learning and attention reveal a general relationship between population activity and behavior. Science 359:463–465.

Oja E (1982) Simplified neuron model as a principal component analyzer. Journal of Mathematical Biology:1–7.

Pouget A, Dayan P, Zemel R (2000) Information processing with population codes. Nature Reviews Neuroscience 1:125–132.

Rabinowitz NC, Goris RL, Cohen M, Simoncelli EP (2015) Attention stabilizes the shared gain of V4 populations. eLife 4:e08998.

Reimer J, McGinley MJ, Liu Y, Rodenkirch C, Wang Q, McCormick DA, Tolias AS (2016) Pupil fluctuations track rapid changes in adrenergic and cholinergic activity in cortex. Nature Communications 7:13289.

Ruff DA, Cohen MR (2014) Attention can either increase or decrease spike count correlations in visual cortex. Nature Publishing Group 17:1591–1597.

Ruff DA, Cohen MR (2016) Stimulus Dependence of Correlated Variability across Cortical Areas. Journal of Neuroscience 36:7546–7556.

Shadlen MN, Newsome WT (1998) The variable discharge of cortical neurons: implications for connectivity, computation, and information coding. J Neurosci 18:3870–3896.

Shamir M, Sompolinsky H (2004) Nonlinear population codes. Neural Comput 16:1105–1136.

Shamir M, Sompolinsky H (2006) Implications of neuronal diversity on population coding. Neural Comput 18:1951–1986.

Smith MA, Jia X, Zandvakili A, Kohn A (2013) Laminar dependence of neuronal correlations in visual cortex. Journal of Neurophysiology 109:940–947.

Stringer C, Michaelos M, Pachitariu M (2019) High precision coding in mouse visual cortex. Biorxiv.

Tsividis P, Pouncy T, Xu JL, Tenenbaum JB, Gershman SJ (2017) Human Learning in Atari. 2017 AAAI Spring Symposium Series, Science of Intelligence: Computational Principles of Natural and Artificial Intelligence:1–4.

Vinck M, Batista-Brito R, Knoblich U, Cardin JA (2015) Arousal and Locomotion Make Distinct Contributions to Cortical Activity Patterns and Visual Encoding. Neuron 86:740–754.

Wimmer RD, Schmitt LI, Davidson TJ, Nakajima M, Deisseroth K, Halassa MM (2015) Thalamic control of sensory selection in divided attention. Nature 526:705–709.

Zanto TP, Gazzaley A (2009) Neural Suppression of Irrelevant Information Underlies Optimal Working Memory Performance. Journal of Neuroscience 29:3059–3066.

Zohary E, Shadlen MN, Newsome WT (1994) Correlated neuronal discharge rate and its implications for psychophysical performance. Nature 370:140–143.

